# Ultrasensitive amplification-free quantification of a methyl CpG-rich cancer biomarker by single-molecule kinetic fingerprinting

**DOI:** 10.1101/2024.04.06.587997

**Authors:** Liuhan Dai, Alexander Johnson-Buck, Peter W. Laird, Muneesh Tewari, Nils G. Walter

## Abstract

The most well-studied epigenetic marker in humans is the 5-methyl modification of cytosine in DNA, which has great potential as a disease biomarker in liquid biopsies of cell-free DNA. Currently, quantification of DNA methylation relies heavily on bisulfite conversion followed by PCR amplification and NGS or microarray analysis. PCR is subject to potential bias in differential amplification of bisulfite-converted methylated *versus* unmethylated sequences. Here, we combine bisulfite conversion with single-molecule kinetic fingerprinting to develop an amplification-free assay for DNA methylation at the branched-chain amino acid transaminase 1 (BCAT1) promoter. Our assay selectively responds to methylated sequences with a limit of detection below 1 fM and a specificity of 99.9999%. Evaluating complex genomic DNA matrices, we reliably distinguish 2-5% DNA methylation at the BCAT1 promoter in whole blood DNA from completely unmethylated whole-genome amplified DNA. Taken together, these results demonstrate the feasibility and sensitivity of our amplification-free, single-molecule quantification approach to improve the early detection of methylated cancer DNA biomarkers.

## Introduction

DNA methylation refers to the addition of a methyl group to a nucleobase—typically to the 5-position of cytosine to form 5-methylcytosine, 5mC—in double-stranded DNA. In mammals, DNA methylation occurs almost exclusively at CpG dinucleotides. Over 70% of CpGs across the human genome are methylated, with the remaining unmethylated CpGs clustered in so-called CpG islands (CGI), regions containing a high proportion of CpGs^1, 2^. Notably, aberrant methylation profiles have been implicated in numerous diseases, and particularly in cancer, where hypermethylation of tumor suppressor gene promoters commonly plays a role in tumorigenesis^1^.

DNA methylation shows promise as a biomarker for the early detection of cancer for several reasons. First, unlike somatic tumor mutation profiles that show significant variation between patients, locus-specific tumor methylation profiles are highly consistent across individuals^1, 3^. Moreover, epigenetic alterations tend to occur early during oncogenesis^4–6^, making them attractive as potential biomarkers for the detection of cancers at an early stage when they are often easier to treat. Additionally, DNA methylation states at specific loci are often tissue-specific and therefore carry information about the tissue of origin of a cancer, which is especially important when measuring tumor-derived DNA in blood (i.e., a liquid biopsy) for cancer diagnosis^7^. Finally, the clustering of concurrent methyl CpGs allows for hybridization-based allele discrimination. Several recent clinical studies showed a strong correlation between hypermethylation in the BCAT1 promoter and tumor progression in colorectal cancer patients^8–14^, and other clinical studies of methylated DNA loci as cancer biomarkers are underway^15, 16^.

Current gold standards for sensitively detecting and/or quantifying DNA methylation rely on amplification via polymerase chain reaction (PCR) or whole-genome amplification following bisulfite conversion, which chemically deaminates unmethylated cytosines to uracils but leaves methylated cytosines intact. These amplification-based approaches—e.g., methylation-specific PCR (MSP)^17^, bisulfite pyrosequencing^18, 19^, bisulfite Illumina sequencing^20–22^, and MethylationEPIC^23, 24^ arrays—generally have low limit of detection^25–28^. However, both overestimation and underestimation of methylation levels at different loci have been suggested due to biased amplification of bisulfite-treated methylated *versus* unmethylated amplicons^24, 29–32^. So far, most efforts to resolve such bias focus on optimization of existing protocols or finding alternatives to bisulfite conversion. However, a relatively unexplored path to reducing bias in quantifying DNA methylation is the use of highly sensitive amplification-free approaches in combination with bisulfite conversion.

Recently, our group has established a sensitive amplification-free detection principle called single-molecule recognition through equilibrium Poisson sampling (SiMREPS)—or single-molecule kinetic fingerprinting—that measures Poisson statistics of individual biomarker molecules by real-time observation of repeated transient interactions with fluorescent probes. Classifying single molecules according to the kinetics and other characteristics of these repeated interactions not only eliminates background signals almost completely but also provides unparalleled specificity when detecting, for example, single nucleotide variants (SNVs). Background-free SiMREPS assays with specificity as high as 99.99998% and sensitivity in the low femtomolar to low attomolar range have been demonstrated for analytes including miRNAs, mutant DNA, and proteins^33–37^. Because it directly detects unamplified single molecules, SiMREPS can avoid PCR bias entirely.

Here we extend the SiMREPS toolbox to a new application: quantification of methylation in the BCAT1 promoter, a biomarker for colorectal cancer^8–14^. We demonstrate 99.9999% discrimination between bisulfite-converted methylated and unmethylated BCAT1 and achieve a limit of detection of 0.368 fM with almost zero background signal. Our assay also shows robust tolerance to high concentrations of unmethylated BCAT1. We further validate our quantification of BCAT1 promoter methylation in a background of genomic DNAs. Our assay reveals a significantly higher level of DNA methylation at the BCAT1 promoter in whole blood DNA from healthy donors (with 2-5% methylation estimated by orthogonal methods) when compared to (unmethylated) whole-genome amplified DNA. The ability of our assay to robustly detect such low levels of DNA methylation in complex genomic DNA matrices suggests potential utility in applications such as early detection of cancer or monitoring of minimal residual disease.

## Results

### General assay pipeline and working principle

Here, we apply the principle of SiMREPS to methylation detection by coupling it with a bisulfite pre-treatment step to yield an assay we term bisulfite methylation-SiMREPS (BSM-SiMREPS). Due to its relevance for cancer diagnostics, we chose a 102 nucleotide (nt) sequence corresponding to a CGI in the BCAT1 promoter to demonstrate BSM-SiMREPS. The target sequence comprises 13 CpG sites (only nine of which are detected and thus shown in **Fig. 1**) whose hypermethylation is linked to colorectal cancer^8, 9, 11, 13, 14, 38^. The assay starts with bisulfite treatment to convert unmethylated cytosines in the double-stranded DNA (dsDNA) target to uracils as shown in **Fig. 1**, introducing mismatches between what were originally fully complementary strands. Each unmethylated cytosine introduces one mismatch, resulting in up to 44 GU mismatches in the bisulfite-converted *methylated* BCAT1 promoter (102 nt MBC) and 70 GU mismatches in the bisulfite-converted *unmethylated* BCAT1 promoter (102 nt UBC). This large number of mismatches ensures that the bisulfite-treated targets will be present in single-stranded form at room temperature (**Supplementary Fig. 1**). Next, two auxiliary probes (Aux1 and Aux2) designed to stably bind the 5mC containing 102 nt MBC—but not the only uracil containing 102 nt UBC—were mixed and heat-annealed with the bisulfite-converted DNAs. Any weak binding of the auxiliary probes to the 102 nt UBC is expected to be completely suppressed in the presence of 2 μM dT_10_ carrier and 10 nM each of sequences Blocker1 and Blocker2, which are complementary to the Aux1 and Aux2 binding sites, respectively, on UBC specifically (see Supplementary Table 1)^34^. The assay mixture was then added to the well of a sample chip precoated with biotin-PEG, streptavidin, and a biotinylated capture probe (CP). As with the auxiliary probes, CP specifically recognizes the 102 nt MBC, and weak interaction with the 102 nt UBC is suppressed by dT_10_^34^. An imaging buffer containing a pair of 8-9 nt fluorescent probes (FPs) labeled with Cy3 and Cy5, respectively, together with an oxygen scavenger system was then added, and the sample well was imaged by total internal reflection fluorescence microscopy (TIRF-M) using an oil-immersion objective.

**Fig. 1.**
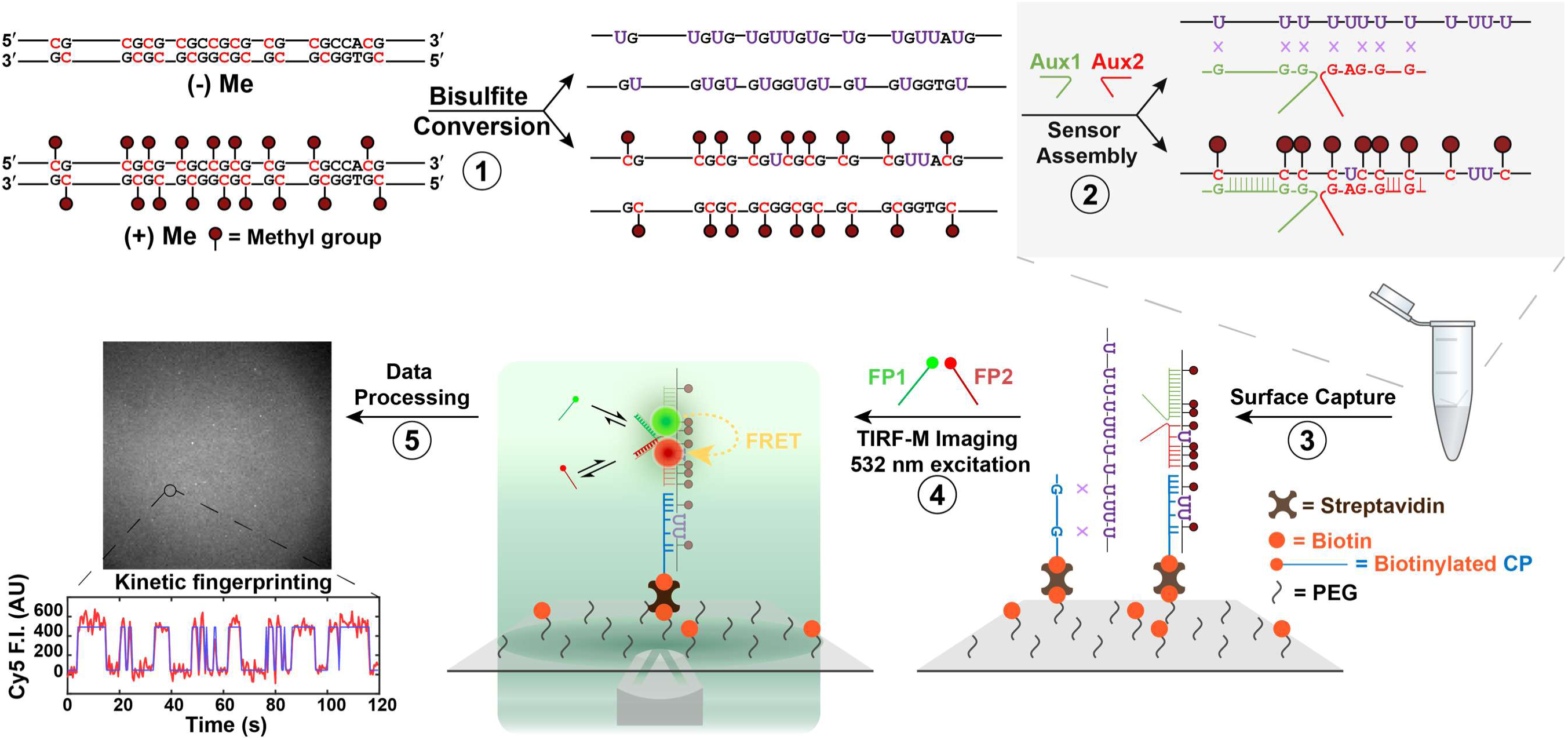
Schematic of the BSM-SiMREPS pipeline. (1) A 102 bp BCAT1 promoter segment (methylated or unmethylated) undergoes bisulfite treatment, converting unmethylated cytosines (C) to uracil (U) and leaving methylated cytosines intact. Up to 44 GU mismatches are introduced as a result, converting the dsDNA fragment to two ssDNAs. (2) The product of bisulfite treatment is mixed with auxiliary probes (Aux1 and Aux2) that bind specifically to the bisulfite-converted, methylated forward strand and provide adjacent docking sites for two fluorescent probes. (3) After probe-target assembly, the mixture is added to a sample well and captured at the surface of a PEGylated coverslip coated with a capture probe *via* the biotin-streptavidin interaction. (4) An imaging buffer containing two fluorescent oligonucleotide probes (FP1 and FP2) is added to the sample well and the surface is illuminated by objective-type TIRF with a 532 nm laser. Cy3 is excited, and Cy5 fluorescence is collected as a FRET readout of simultaneous association of FP1 and FP2 to the same target molecule. (5) During data processing, individual fluorescent spots (candidate molecules) are identified and converted to intensity-time traces which are analyzed with a hidden Markov model to identify “on” and “off” states. Kinetics and other signal characteristics are used to distinguish genuine target signals from background signals in a process known as kinetic fingerprinting. F.I., fluorescence intensity; AU, arbitrary unit.

Simultaneous binding of both FPs to the docking sites on their respective auxiliary probes on the same target molecule results in Förster resonance energy transfer (FRET) from the Cy3-labeled FP to the Cy5-labeled FP, yielding a detectable Cy5 fluorescence signal under excitation of Cy3 (**Fig. 1**). Thus, repeated simultaneous occupancy of a target by both FPs yields an alternating pattern of high and low Cy5 fluorescence over time, yielding a distinct kinetic fingerprint that permits identification and quantification of immobilized sensor molecules. Background signal is reduced to nearly zero in BSM-SiMREPS by three mechanisms: 1) background fluorescence originating from FPs in solution is minimized by the restricted excitation and detection volume of TIRF-M; 2) fluorescence from FPs entering the TIRF excitation volume is minimized by FRET detection, since two FPs are unlikely to be within FRET distance of one another in absence of interactions with a common target molecule; 3) any remaining signal bright enough to be detectable is unlikely to exhibit kinetic characteristics similar to those arising from repeated FP interaction with a genuine target, and can thus be rejected by applying kinetic filtering criteria.

### Initial optimization of assay design and FP pair sequences

To decouple FP design from complications such as target secondary structure and chemical damage by bisulfite, we first employed a series of shorter and mimic versions of the bisulfite-converted methylated BCAT1 promoter. The target mimics are unmethylated DNAs directly synthesized to reflect the expected sequences of bisulfite-converted products and do not undergo bisulfite treatment (see Supplementary Table 1 and **Fig. 2a**). Initially, we tested a 42 nt MBC Mimic (**Fig. 2b**, 42 nt Construct) with a scheme involving two probes: Aux1, which provides a docking site for FP1, and Biotin-Aux2, a biotinylated second auxiliary probe (see Supplementary Table 1 and **Fig. 2b**, 42 nt Construct), which serves as a capture probe and provides a docking site for FP2 (**Fig. 1**). However, this construct resulted in high background signal and poor signal-to-noise (S/N) ratio in kinetic fingerprinting (**Fig. 2c**, 42 nt Construct). We hypothesized that this high background was caused by a high surface density of Biotin-Aux2, which may recruit FP2 to the surface regardless of the presence or absence of target. With several thousand or even millions of copies of FP2 bound to the surface within a field of view, even a small amount of direct excitation of the Cy5 acceptor at 532 nm could yield high background signal. To circumvent this problem, we designed a new 55 nt MBC Mimic construct (see Supplementary Table 1, **Fig. 2b**, 55 nt Construct) incorporating a third, dedicated biotinylated capture probe (CP1) that recruits the target to the surface independently of the two auxiliary probes. As predicted, TIRF-M imaging of the 55 nt MBC Construct revealed much lower background signal, permitting us to easily detect bright fluorescent spots arising from FRET between transiently binding FP1 and FP2, and to analyze the resulting kinetic fingerprints with high S/N (**Fig. 2c**, 55 nt Construct).

**Fig. 2.**
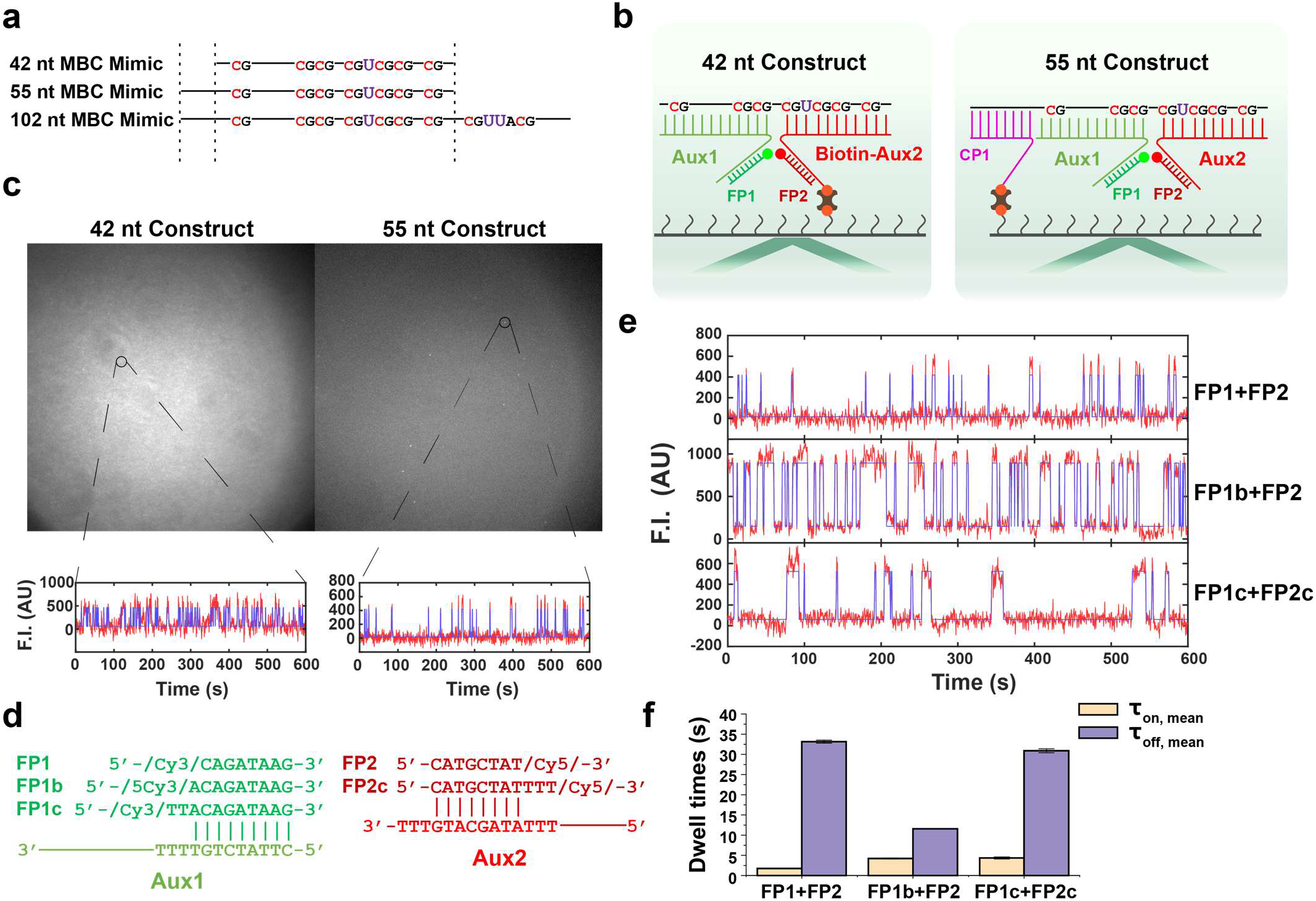
Optimization of assay constructs and fluorescent probe sequences. **a** Sequence-aligned schematics of MBC Mimics of different lengths showing positions of CpG dinucleotides and uracil bases introduced to mimic bisulfite-converted unmethylated cytosines. **b** In the 42 nt construct (designed for detection of 42 nt MBC Mimic), Aux2 serves both as an auxiliary probe and a biotinylated capture probe. In the 55 nt construct (designed for detection of 55 nt MBC Mimic), a separate biotinylated capture probe CP1 is introduced along with the two auxiliary probes to decouple surface capture from FP recruitment. **c** Raw TIRF microscopy video frames of assays conducted with the 42 and 55 nt constructs (top) and representative intensity-time traces (bottom, red lines) fit by HMM (blue lines). **d** Sequences of different FPs and their binding sites on Aux1 or Aux2. **e** Representative intensity-time traces (red lines) fit by HMM (blue lines) for different FP pairs. **f** Comparison of mean *τ*_*on*_ and *τ*_*off*_ of each pair of imagers. Mean *τ*_*on*_ and *τ*_*off*_ are calculated by fitting a single exponential decay function to a cumulative histogram of dwell times of individual high- or low-fluorescence events in all traces. Error bars represent standard errors of fitted parameters, calculated as the square root of variance estimate for the parameter. F.I., fluorescence intensity; AU, arbitrary unit.

Using the 55 nt MBC Construct, different FP pairs were tested to optimize kinetics for rapid, high-confidence detection of the target sequence (**Fig. 2d,f**). The initial probe pair FP1 + FP2 exhibited long dwell times in the unbound state (*τ*_*off*_) and very short dwell times in the bound state (*τ*_*on*_). In contrast, the pair FP1b + FP2—wherein one AT base pair is added between FP1b and Aux1—yielded more similar *τ*_*on*_ and *τ*_*off*_ values, which is desirable as it maximizes the average number of signal transitions in a fixed observation window^36, 39^. In an attempt to increase FRET efficiency, we also tested the combination FP1c + FP2c, wherein 2T and 3T linkers are added at the fluorophore-labeled end of each FP, placing the fluorophores closer together in theory (**Fig. 2d**). However, this significantly increased *τ*_*off*_ while leaving *τ*_*on*_ unchanged (**Fig. 2f**), suggesting that the added linkers may introduce steric hindrance that reduces the binding rate constant *k*_*on*_. Since the combination FP1b + FP2 showed the least bias between *τ*_*on*_ and *τ*_*off*_, we chose this pair of FPs for use in the assay.

Next, we further optimized imaging conditions including temperature and FP concentration to shorten acquisition time while maintaining analytical performance (**Supplementary Fig. 2** and Supplementary Note 1). Different CPs were also evaluated and CP2 was chosen because it yielded the best combination of assay sensitivity, reproducibility, and low false positives in the presence of the 102 nt UBC Mimic (**Supplementary Fig. 3** and Supplementary Note 1).

### Detection of mimic and non-mimic targets with BSM-SiMREPS

The final assay design employs Aux1, Aux2 and CP2 to selectively immobilize methylated BCAT1 promoter, followed by imaging with 100 nM FP1 and 100 nM FP2 for 2 min at 26.5 °C to selectively detect the methylated target (**Fig. 3**): i.e., 102 nt MBC or MBC Mimic (**Fig. 3a**). To assess the performance of our assay with both mimic and non-mimic targets, we conducted the assay in the presence of either 1 pM 102 nt MBC or MBC Mimic as methylation-positive controls, either 5 nM 102 nt UBC or UBC Mimic as methylation-negative controls, and a no-target (blank) control (**Fig. 3b-d**). Measurements revealed clearly distinct kinetic fingerprints between methylation-positive and methylation-negative controls in the case of both mimic and non-mimic targets (**Fig. 3b**). Kinetic fingerprints of methylation-positive target molecules can be distinguished from those of non-target signals and methylation-negative molecules on the basis of the number of binding and dissociation events per molecule (*N*_*b*+*d*_, **Fig. 3c**) and the median values of dwell times *τ*_*on*_ and *τ*_*off*_ (**Fig. 3d**). Interestingly, the *N*_*b*+*d*_ distribution of the 102 nt MBC is shifted to the left compared to that of the 102 nt MBC Mimic due to a slightly increased *τ*_*off*_ (**Fig. 3c** and **Supplementary Fig. 4a**). One possible explanation is that DNA damage such as depurination or depyrimidination after bisulfite treatment reduces the stability of auxiliary probe assembly. MBC molecules containing abasic sites might still interact with the auxiliary and capture probes, but the auxiliary probes could partially and reversibly dissociate from these MBC molecules, separating the donor and acceptor fluorophores sufficiently so that FRET no longer takes place, resulting in longer low-FRET dwell times and a larger median *τ*_*off*_. Partial dissociation of auxiliary probes would also be predicted to lower the average FRET efficiency between bound pairs of FPs; consistent with this prediction, we observed a significant decrease in “on” state fluorescence intensity when detecting bisulfite-converted MBC rather than the MBC Mimic (**Supplementary Fig. 4b**). Notably, we did not observe any evidence of dissociation of the auxiliary probes from the MBC or MBC Mimic by Native PAGE (**Supplementary Fig. 4c**), suggesting that the probes remain largely bound to both targets and any dissociation is either partial or short-lived and hence might be difficult to visualize on a gel. In any case, although the kinetic fingerprints of the MBC and MBC Mimic are similar, the subtle differences between them highlight the importance of calibrating SiMREPS assays with reference standards that closely resemble the intended targets.

**Fig. 3.**
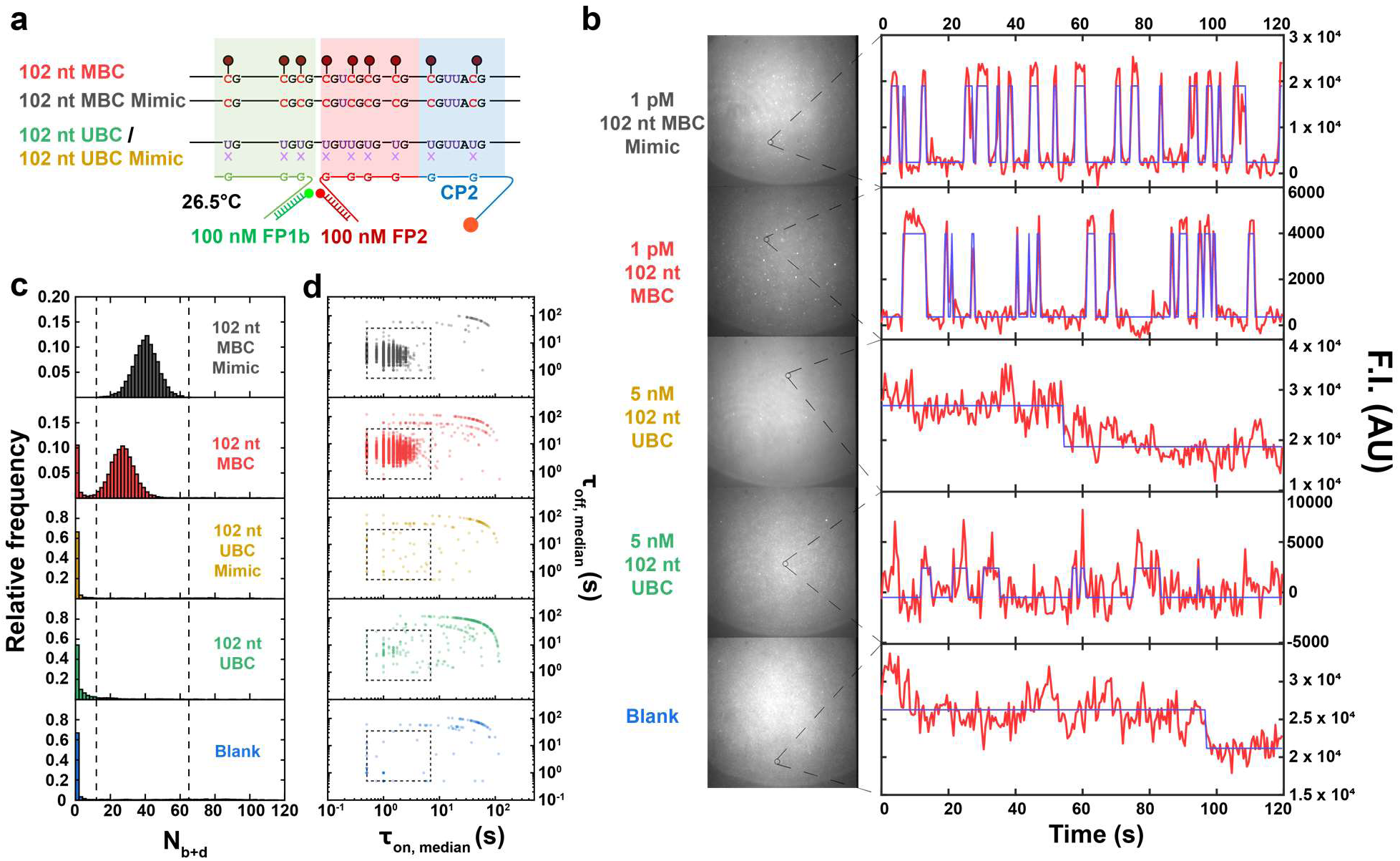
Detection of different types of samples using BSM-SiMREPS. **a** Sequence-aligned schematics of the forward strands of 102 nt methylation-positive (MBC, MBC Mimic) and methylation-negative (UBC, UBC Mimic) targets and target mimics. The 102 nt MBC Mimic shares the same sequence as the 102 nt MBC but without methyl groups and without the accompanying bisulfite-converted methylated reverse strand. The 102 nt UBC Mimic shares the same sequence as 102 nt UBC but without the accompanying bisulfite-converted unmethylated reverse strand. Shaded regions represent binding sites for auxiliary and capture probes. **b** Raw TIRF microscopy video frames from assays of different target, target-mimic, or blank conditions and corresponding representative intensity-time traces (red lines) fit by HMM (blue lines). **c** Distribution of *N*_*b*+*d*_ derived from SiMREPS analysis of each sample type. Dashed lines represent thresholds used to distinguish MBC and MBC Mimic from UBC, UBC Mimic, and blank conditions. **d** Distribution of median *τ*_*on*_ and *τ*_*off*_ of each trace in each sample type. Dashed lines represent thresholds used to distinguish MBC and MBC Mimic from UBC, UBC Mimic, and blank conditions.

### Sensitivity and specificity of BSM-SiMREPS assay

Next, we evaluated the sensitivity and specificity of BSM-SiMREPS for the methylated BCAT1 promoter by assaying both the MBC and MBC Mimic target at varying concentrations, and in the presence or absence of non-target UBC or UBC Mimic (**Fig. 4**). Kinetic filtering criteria were optimized based on positive and negative control measurements, and used to identify and count the number of target molecules (i.e., accepted counts) within each field of view (FOV). For improved sensitivity, we combined the total accepted counts from 10 FOVs (**Supplementary Fig. 5**)^37^. Assays of both 1 pM MBC Mimic and 1 pM MBC showed similar signal levels of ∼10,000 total accepted counts, with an average of <1 accepted count in blank measurements (**Fig. 4a**). Consistent with the expected high specificity of BSM-SiMREPS, even a 5,000-fold higher concentration (5 nM) of the UBC or UBC Mimic yielded approximately 200- to 2,000-fold lower signal than 1 pM of MBC or MBC Mimic, implying a specificity of ≥99.9999%. Despite the subtle differences in kinetic fingerprints discussed above, the standard curves of the MBC Mimic and MBC targets were nearly identical in slope and y-intercept, yielding estimated limits of detection (LODs) of 0.365 fM and 0.368 fM, respectively (**Fig. 4b**). Furthermore, standard curves collected in the presence of a constant background of 1 nM (=1,000 pM) UBC Mimic or UBC (**Fig. 4c**) maintained good linearity for both targets and nearly the same LOD as for the MBC target; however, the LOD for the MBC Mimic increased 4.6-fold, suggesting some interference from the UBC Mimic.

**Fig. 4.**
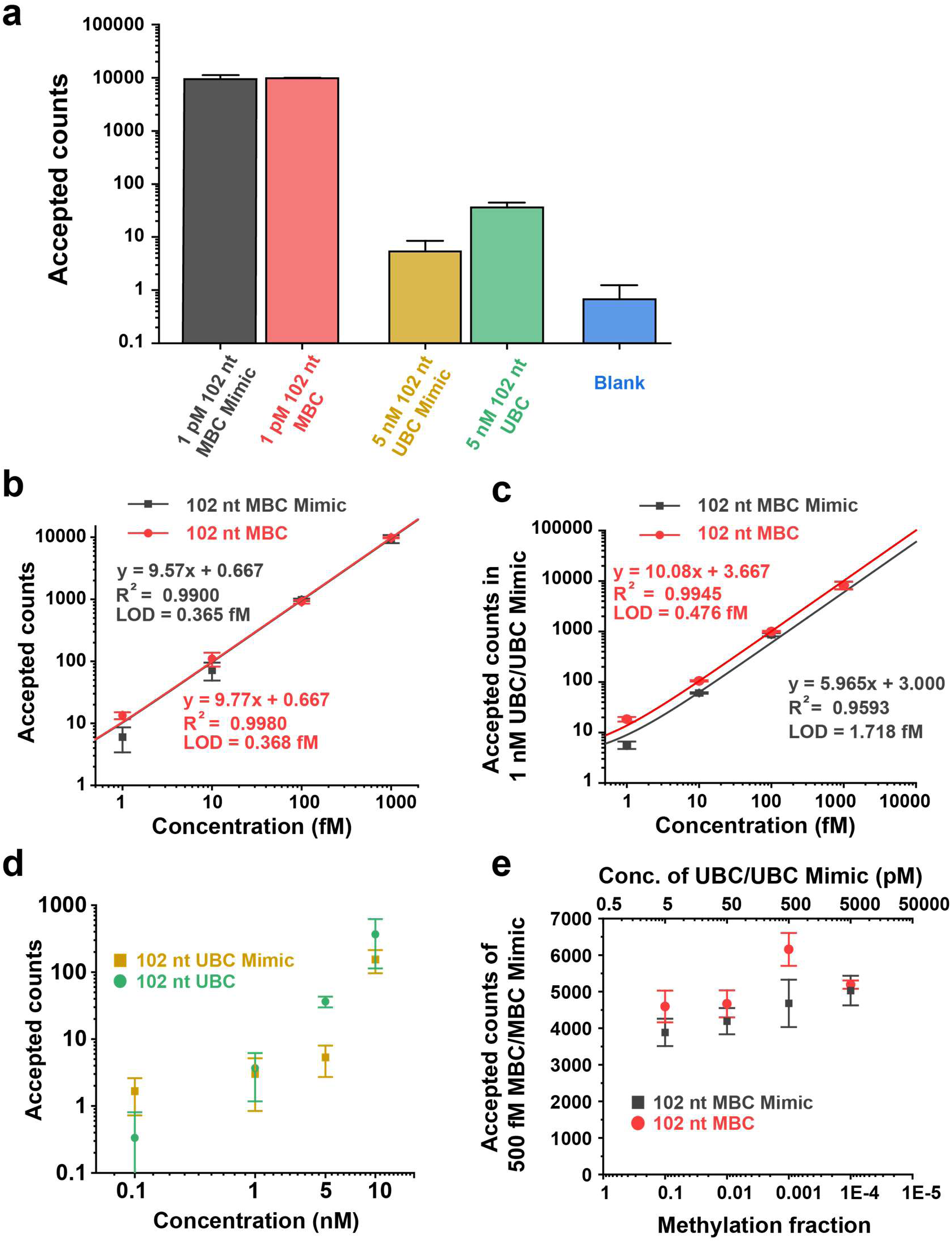
Quantification of MBC and MBC Mimic and assessment of analytical performance. **a** Mean accepted counts from BSM-SiMREPS assays of 1 pM methylated target or mimic, 5 nM (5000 pM) unmethylated target or mimic, or a blank sample. **b** Standard curves from BSM-SiMREPS assays of 102 nt MBC Mimic and 102 nt MBC. **c** Standard curves from BSM-SiMREPS assays of 102 nt MBC Mimic in a background of 1 nM 102 nt UBC Mimic and of 102 nt MBC in a background of 1 nM 102 nt UBC. **d** False positive counts measured at varying concentrations of 102 nt UBC Mimic and UBC. **e** Accepted counts from BSM-SiMREPS assays of 500 fM 102 nt MBC in the presence of different concentrations of 102 nt UBC and of 500 fM 102 nt MBC Mimic in different concentrations of 102 nt UBC Mimic. Datapoints in panel a-f are presented as mean ± 1 s.d., with n = 3 independent experiments. Accepted counts represent the sum over ten FOVs for each condition.

To further characterize background signal from unmethylated targets, we assessed the number of false positives in varying concentrations of UBC Mimic or UBC without any methylated promoter present (**Fig. 4d**). At 10 nM UBC Mimic or UBC, accepted counts were unstable, decreasing over time as sequential FOVs were measured (**Supplementary Fig. 6**), perhaps due to dissociation of weakly captured UBC or UBC Mimic from the surface. Therefore, we considered 5 nM as the upper limit for measuring specificity since accepted counts were consistent across sequential FOVs. Next, we measured the robustness of our assay in detecting 500 fM MBC Mimic or MBC in varying concentrations of UBC Mimic or UBC (**Fig. 4e**). A slight increase (∼15-30%) in accepted counts was observed in assays of both mimic and non-mimic target as the concentration of UBC Mimic or UBC increased to a ∼10,000-fold excess relative to methylated target (**Fig. 4e**). Surprisingly, accepted counts for the MBC target spiked at 500 pM UBC, corresponding to a methylation fraction of 0.1%. No significant difference in kinetic fingerprinting was observed as a function of UBC concentration. One possible explanation for this peak in signal is that 500 pM of UBC improves surface capture of MBC by reducing non-specific absorption of MBC to the surface or the wall of the sample well; at higher concentrations, UBC might begin to compete with MBC for capture probes, countering this effect.

Compiling results across experimental conditions (**Table 1**) and estimating both discrimination factor and specificity as described elsewhere^36, 40^, we find that BSM-SiMREPS consistently detects hypermethylated BCAT promoter with a specificity of ∼99.9999%.

**Table 1.**
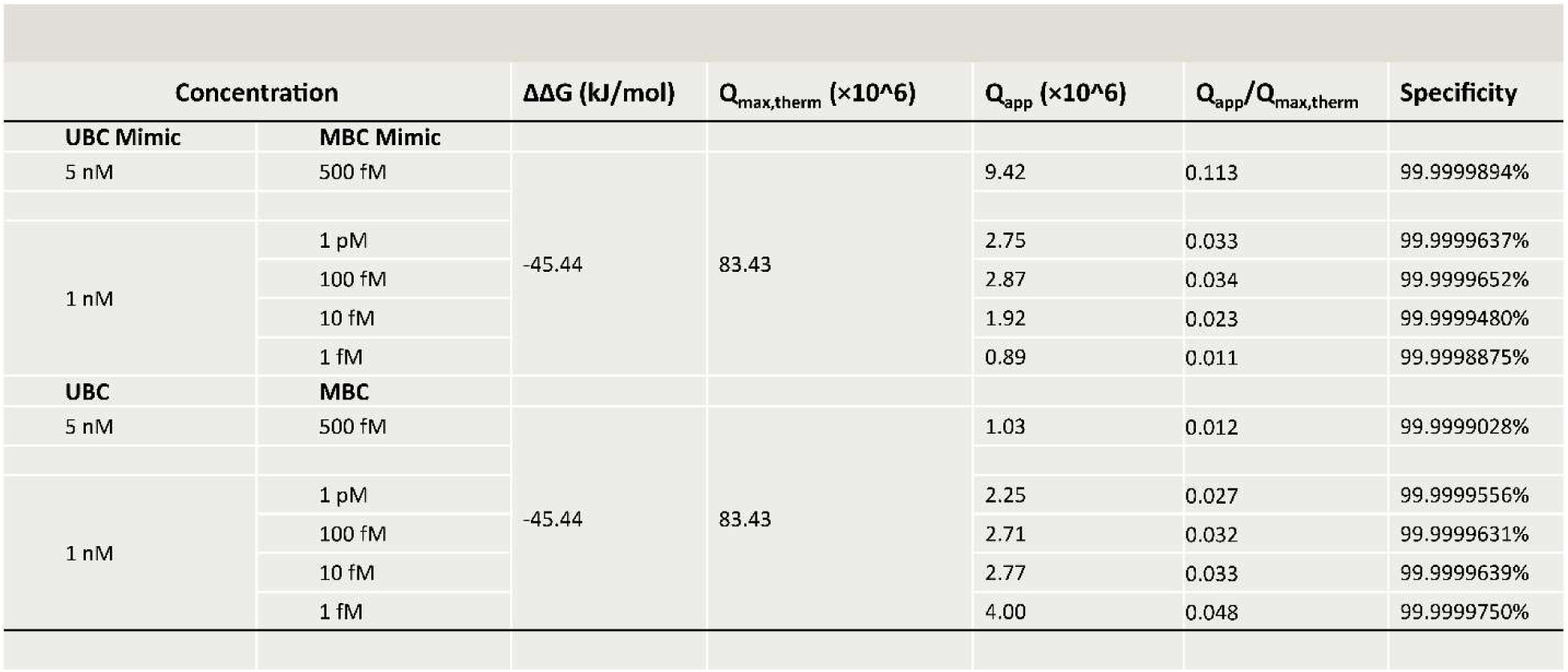
Maximum thermodynamic discrimination factor (Q_max,therm_) as well as apparent specificity and discrimination factors (Q_app_) calculated at different experiment conditions.

| Concentration | | $\Delta\Delta G$ (kJ/mol) | $Q_{\max, \text{therm}}$ ($\times 10^6$) | $Q_{\text{app}}$ ($\times 10^6$) | $Q_{\text{app}}/Q_{\max, \text{therm}}$ | Specificity |
| --- | --- | --- | --- | --- | --- | --- |
| **UBC Mimic** | **MBC Mimic** |  |  |  |  |  |
| 5 nM | 500 fM | -45.44 | 83.43 | 9.42 | 0.113 | 99.9999894% |
| 1 nM | 1 pM |  |  | 2.75 | 0.033 | 99.9999637% |
|  | 100 fM |  |  | 2.87 | 0.034 | 99.9999652% |
|  | 10 fM |  |  | 1.92 | 0.023 | 99.9999480% |
|  | 1 fM |  |  | 0.89 | 0.011 | 99.9998875% |
| **UBC** | **MBC** |  |  |  |  |  |
| 5 nM | 500 fM | -45.44 | 83.43 | 1.03 | 0.012 | 99.9999028% |
| 1 nM | 1 pM |  |  | 2.25 | 0.027 | 99.9999556% |
|  | 100 fM |  |  | 2.71 | 0.032 | 99.9999631% |
|  | 10 fM |  |  | 2.77 | 0.033 | 99.9999639% |
|  | 1 fM |  |  | 4.00 | 0.048 | 99.9999750% |

### Detection of BCAT1 promoter methylation in a background of genomic DNA

Finally, we evaluated the BSM-SiMREPS assay’s ability to quantify methylated BCAT1 promoter in a background of genomic DNA. To do so, we spiked varying concentrations of 102 nt MBC into two different bisulfite-converted genomic DNA matrices: an absolute methylation-negative control^41^–whole-genome amplified DNA (BS WGA) from DKO (double-knock-out) HCT116 cell lines ([DNMT1 (-/-) / DNMT3b (-/-)]), and DNA extracted from human male whole blood (BS Blood DNA) (**Fig. 5**, **Supplementary Table 1**). In the presence of genomic DNA, we observed higher background fluorescence, presumably due to non-specific interaction of FPs with surface-adsorbed genomic DNA fragments (**Fig. 5a**). Nevertheless, single molecules of 102 nt MBC were still easily distinguished from this background due to their distinctive kinetic fingerprints. Interestingly, we detected significant BCAT1 methylation in BS Blood DNA but not in BS WGA (**Fig. 5b**). Importantly, 20 fM haploid bisulfite-converted fully methylated whole-genome amplified DNA from DKO HCT116 (BS +Me WGA, see **Supplementary Table 1**) yielded nearly identical accepted counts as 20 fM MBC spiked into a background of 20 fM haploid BS WGA, indicating a spike-in recovery of ∼100% for BSM-SiMREPS in this matrix (**Fig. 5b**). Next, we constructed calibration curves for MBC in both BS Blood DNA and BS WGA (**Fig. 5c**), resulting in estimated LODs of 1.33 fM and 1.62 fM, respectively. The slight increase in these LODs compared to that seen in a background of 102 nt UBC might arise from two factors: compromised capture efficiency due to competition from genomic DNA, as well as the increased background signal discussed above. Consistent with our prior observation of methylation-positive signal in BS Blood DNA lacking spike-in, we observed a constant offset of ∼10 accepted counts in BS Blood DNA compared to BS WGA at each concentration of 102 nt MBC (**Fig. 5c**).

**Fig. 5.**
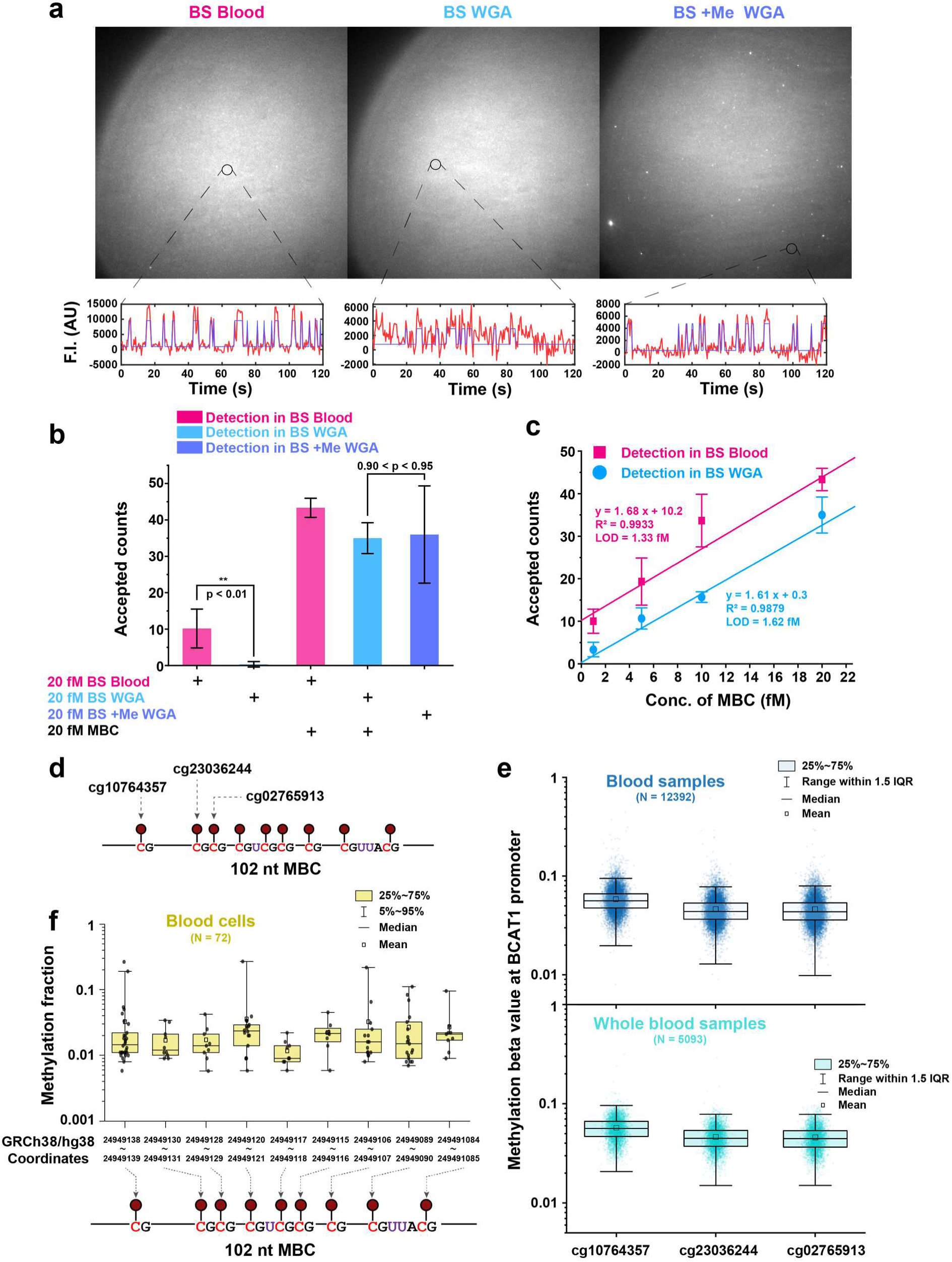
BSM-SiMREPS quantification of 102 nt MBC in a background of two types of genomic DNAs and comparison with bisulfite sequencing and Illumina Infinium MethylationEPIC microarray (EPIC array) at the BCAT1 promoter. **a** Raw TIRF microscopy video frames for BSM-SiMREPS assays of three different types of genomic DNA samples and corresponding representative intensity-time traces (red lines) fit by HMM (blue lines). **b** Quantification of methylated BCAT1 promoter in different genomic DNA samples at 20 fM haploid concentration, with P-value as assessed using a single-tailed, unpaired t-test. **c** Standard curves for 102 nt MBC spiked into two types of genomic DNA samples. **d** Illustration of three cg probes covering BCAT1 promoter used in EPIC array (GEO accession ID: GPL21145). **e** Distributions of methylation beta values at the BCAT1 promoter region for all blood samples, as well as the subset comprising whole blood samples specifically, using the EPIC array. Data comprise all studies on GEO as of December 2022 and were compiled using recountmethylation. **f** Distributions of methylation fractions using bisulfite-sequencing at each CpG site in the BCAT1 promoter in blood cells.

To better interpret our detection of methylated BCAT promoter in whole blood, we first collected the results of existing microarray-based assays and bisulfite sequencing assays in blood samples (**Fig. 5d-f**) and, second, validated these results using bisulfite pyrosequencing on the same batch of whole blood DNA that had been tested by BSM-SiMREPS (**Supplementary Fig. 8**). Maden *et al.* published an R/Bioconductor package called “recountmethylation,” allowing easy access to preprocessed methylation beta values measured by the Illumina Infinium MethylationEPIC microarray (EPIC array). Using recountmethylation, we compiled methylation beta values at the BCAT1 promoter from 12,392 samples on blood samples available in GEO^42^. On the EPIC array, three degenerate cg probes are available within the 102 nt BCAT1 promoter (**Fig. 5d**). Distributions of methylation beta values measured in all blood samples, as well as the subset consisting specifically of whole blood samples, are shown in **Fig. 5e**. The ranges of beta values seen for all three probes are comparable with each other, with median values of approximately 5%. Turning to existing bisulfite sequencing data, we manually combined 72 tracks measured for different subtypes of blood cells at the BCAT1 promoter from bisulfite sequencing studies on the UCSC genome browser^43^. The bisulfite sequencing data show methylation level distributions with median values of approximately 2% (although the ranges of some CpG sites extend to 10-30%). Our targeted bisulfite pyrosequencing similarly measured a mean value of 1.7% methylation at the BCAT1 promoter in our whole blood DNA sample (**Supplementary Fig. 8b**), consistent with bisulfite sequencing results in other whole blood samples (**Fig. 5d**). Altogether, these results consolidate a 2-5% methylation level at the BCAT1 promoter in whole blood DNA. Notably, targeted bisulfite pyrosequencing can only discriminate BS Blood DNA *versus* BS WGA with a P-value of higher than 0.10 (**Supplementary Fig. 8b**), whereas our assay measured significantly higher level in BS Blood compared to BS WGA, with a P-value lower than 0.01 (**Fig. 5b)**. Our assay also showed nearly zero counts in BS WGA (**Fig. 5b)**, whereas targeted bisulfite pyrosequencing presented a basal level of DNA methylation with a relatively broad distribution across 0.7% up to 4.1% (**Supplementary Fig. 8b)**. This subtle difference in DNA methylation level between BS Blood DNA and BS WGA might be lost during PCR amplification but can be reliably detected in our assay, presumably due to the amplification-free nature of BSM-SiMREPS.

## Discussion

In this study, we developed a methylation-specific single-molecule fluorescence kinetic fingerprinting assay by combining bisulfite treatment with SiMREPS, and applied it to the detection of hypermethylation in the BCAT1 promoter, a biomarker associated with colorectal cancer. By employing TIRF, FRET, and careful probe design, we achieved high signal-to-noise detection of repeated FP binding to individual target-probe complexes. Optimization of FP pair sequences, concentrations, and imaging temperature resulted in an ultra-low background assay whose sensitivity was increased further by imaging multiple FOVs per sample well. Ultimately, we achieved a sub-femtomolar LOD and a specificity of 99.9999% for methylated over unmethylated BCAT1 promoter. Furthermore, we demonstrated amplification-free, direct quantification of BCAT1 promoter methylation in the presence of genomic DNA from different sources. Finally, in applying the assay to endogenous BCAT1 promoter in human whole blood DNA, we found a significant fraction of methylated BCAT1 compared to whole-genome amplified DNA, with a signal of ∼10 counts equivalent to 2-5% methylation level. These results show our assay’s ability to detect low levels of DNA methylation even in complex genomic DNA matrices. Compared to our assay, targeted bisulfite pyrosequencing showed lower statistical significance in discriminating minimal DNA methylation at the BCAT1 promoter in whole blood DNA *versus* whole-genome amplified DNA, highlighting the promise of our assay as a better candidate for early detection of methylation cancer biomarkers.

Several issues arose in bisulfite conversion. One was bias in the yield of forward and reverse strands of Me-BCAT1 following bisulfite conversion that arose when certain lots of the bisulfite conversion kit were used (see **Supplementary Fig. 9**). However, we did not observe such bias when bisulfite converting WGA since spike-in recovery is ∼100% as previously discussed (**Fig. 5b**). We therefore bisulfite-converted Me-BCAT1 Forward and Reverse individually and then mixed them in an equimolar ratio to compensate for biased yield during bisulfite treatment. Other issues include loss of material, DNA damage, and reaction selectivity^44–46^. Both incomplete conversion of unmethylated cytosines and over-conversion of methylated cytosines have the potential to compromise capture efficiency and assay specificity^46–48^.

Our current LOD is still limited by capture efficiency and detection efficiency^49^ and thus a high concentration of extracted genomic DNAs (20 fM or ∼40 ng/µl) is still required. Capture efficiency is the percentage of target molecules in a sample well that are immobilized on the surface, while detection efficiency is the percentage of surface-immobilized molecules that are detected by the assay. Given the field of view of our 60× TIRF objective and the physical size of our camera chip, one FOV in the sample plane is 136.53 µm × 136.53 µm in size. Since our sample wells have an inner diameter of 5.842 mm, the total area imaged in our assay (10 FOVs) comprises only about 0.7% of the capture surface and, hence, only about 0.7% of captured target molecules. Furthermore, diffusion-limited mass transport typically yields a capture efficiency of only 0.5% to 1.5% in similar surface-based assays^35^. Taking both capture and detection efficiency into account, the total analytical efficiency is only 3.5 to 10.5^10^-5^. Thus, if a larger proportion of molecules can be captured and/or detected using pre-enrichment methods like aqueous-two-phase systems^37^, there is potential to increase the sensitivity of BSM-SiMREPS to the low-attomolar or zeptomolar range.

## Methods

### Oligonucleotides

All 102 nt targets were purchased from Integrated DNA Technologies (IDT, www.idtdna.com) with Ultramer oligo synthesis and standard desalting purification. All other DNA oligonucleotides were purchased from IDT with standard synthesis and desalting purification. All fluorescent probes (FPs) with either 5′ Cy3 or 3′ Cy5 modifications were purchased from IDT with high-performance liquid chromatography (HPLC) purification. The target sequence resides in the promoter of BCAT1, with genomic coordinates Chr12: 24,949,058 – 24,949,159 (genome build: UCSC Genome browser GRCh38/hg38 version)^8^; sequences of truncated 55- and 42-nt model targets are shown in **Supplementary Table 1**. All mimics of the methylated target have the same sequence as the corresponding segment of the bisulfite-converted methylated BCAT1 promoter but are unmethylated. Similarly, all mimics of the non-methylated target have the same sequence as the corresponding segment of the bisulfite-converted unmethylated BCAT1 promoter. WGA, +Me WGA, and Blood DNA were obtained from commercial vendors as indicated in **Supplementary Table 1**. See **Supplementary Table 1** for names, sequences, and descriptions of all oligonucleotides, including targets and target mimics.

### Bisulfite treatment

Phosphate buffered saline (PBS) was either diluted from 10× PBS stock solution (Gibco™ PBS (10×), pH 7.4, Fisher Scientific, Cat. No. 70-011-044) using Invitrogen™ UltraPure™ DNase/RNase-Free Distilled Water (ThermoFisher, Cat. No. 10977015) or directly purchased from Fisher Scientific (Gibco™ PBS, pH 7.4, Fisher Scientific, Cat. No. 10-010-023). Double-stranded DNA substrates were prepared by combining complementary single-stranded oligonucleotides at ∼1 μM final concentration of each oligonucleotide in 4× PBS (4× phosphate-buffered saline: 40 mM Na_2_HPO_4_, 7.2 mM KH_2_PO_4_, pH 7.4, 548 mM NaCl, 10.8 mM KCl), heating at 90 °C for 5 min, cooling to 37 °C for 5 min, and finally holding at room temperature for 10 min before storing at −20 °C for further use. Bisulfite treatment was conducted using the EZ DNA Methylation-Lightning^TM^ Kit (Zymo Research, Cat. No. D5030) according to the manufacturer’s protocol. Briefly, DNA substrates were mixed with Lightning Conversion Reagent were first heated at 98 °C for 8 min, annealed to 54 °C for 1 h, and finally held at 4 °C before desulfonation. Desulfonation was carried out for 15 min followed by column purification. The product was eluted into a volume of ∼20 μL. Concentrations of eluted bisulfite-converted genomic DNAs were determined by Nanodrop (NanoDrop 2000, Thermofisher, Cat. No. ND-2000) with 1 A260 = 33 ng/µL as of ssDNA. Concentrations of eluted DNA other than genomic DNAs were determined by Qubit (Qubit™ ssDNA Assay Kit, ThermoFisher, Cat. No. Q10212) from 3 independent measurements.

Each 150-μL bisulfite conversion reaction only accommodates a maximum of 600 ng genomic DNAs at most to ensure optimal conversion efficiency and specificity. In particular, we observed overloading of genomic DNAs to cause significant false positives from incomplete conversion (data not shown). To accommodate larger quantities of genomic DNA (Blood DNA, WGA, and +Me WGA), we therefore performed multiple bisulfite conversion reactions in parallel for each sample type and, following elution from the purification columns, eluted product from individual reactions were combined and concentrated by vacuum centrifugation.

### Design of BSM-SiMREPS probes

Auxiliary probes and capture probes were designed to stably bind target sequences under BSM-SiMREPS assay conditions. We considered three main factors in designing their sequences: 1) melting temperatures (*T*_*m*_) of their hybridization with the target should be ≥ 10 °C above the capture and imaging temperature; 2) upon hybridization with target, binding regions of two neighboring strands should be at least 3 nt away on the target sequence to avoid steric hindrance, but still well within the Förster radius of ∼5 nm; 3) the 5′ end of the capture probe should be biotinylated to ensure that any truncated synthesis products (which might not stably capture the target) are not immobilized on the surface. The *T_m_* of interactions between auxiliary probes or capture probes and the target was estimated using the NUPACK web application (http://www.nupack.org/partition/new) using the following parameters: number of strand species = 2, maximum complex size = 2, minimum temperature = 10 °C, increment = 3 °C, maximum temperature = 70 °C, target DNA concentration = 1 pM, probe concentration = 1 pM, [Na^+^] = 646 mM. From the NUPACK-generated melting curve, we computed the first derivative of the fraction of unpaired bases and estimated the *T_m_* as the temperature at which this first derivative is maximal (see Supplementary Note 3, **Supplementary Fig. 11**).

FP sequences were designed as follows. Unlike those of auxiliary and capture probes, FP sequences are not constrained by the target sequence, but we considered the following factors in their design: 1) no secondary structure should be predicted by NUPACK under assay conditions; 2) no self-hybridization (e.g., dimerization) should be predicted by NUPACK under assay conditions; 3) no interaction longer than 4 base pairs should be predicted to any exposed sequence of the capture probes minimize the potential for significant background signal; and 4) the *T_m_* of interactions of FPs with their docking sites on auxiliary probes should be close to room temperature under assay conditions. *T_m_* values were estimated using NUPACK as described above, but using the following parameters: minimum temperature = 0 °C, maximum temperature = 50 °C, auxiliary probe concentration = 1 nM, fluorescent probe concentration = 100 nM (**Supplementary Fig. 11**).

### BSM-SiMREPS assays

Sample cells made of cut P20 pipette barrier tips were attached to glass coverslips passivated with a 1:10 mixture of biotin-PEG and mPEG. A detailed protocol of slide preparation is discussed elsewhere^36, 39^. Sample cells were first washed with T50 buffer (10 mM Tris-HCl [pH 8.0 at 25 °C], 50 mM NaCl) and then incubated with 40 μL 0.25 mg/mL streptavidin in T50 buffer for 10 min. Following three washes with 1× PBS, the well was incubated for 10 min with a solution of 100 nM capture probe in 1× PBS that had been preheated at 90 °C for 5 min in a metal bath, annealed at 37 °C for 5 min in a water bath, and cooled down to room temperature before addition to the sample well. The sample well was washed three times with 4× PBS, and the last wash left in until addition of the target mixture. The target mixture was prepared by adding the target strand (bisulfite-converted methylated or unmethylated double-stranded BCAT1 promoter or a single-stranded mimic) to a PCR tube containing a final concentration of 10 nM each of Aux1, Aux2, blocker 1, and blocker 2 in 4× PBS with 2 μM poly-T oligodeoxyribonucleotide (dT_10_) as a carrier. All dilutions of targets were performed in 4× PBS with 2 μM dT_10_ in GeneMate low-adhesion 1.7-mL microcentrifuge tubes (VWR, Cat No. 490003-230). PCR tubes containing target mixtures were then heated in a thermocycler to 73 °C for 3 min, annealed at 46.6 °C for 5 min followed by 40 °C for another 5 min, and finally cooled to 25 °C. The annealed target mixtures were then added to the capture probe-coated sample cell and incubated for 1 h at room temperature. Next, sample cells were washed 3 times with 4× PBS, followed by addition of 100 µL imaging buffer containing 100-300 nM (in the final assay, 100 nM) of each FP in the presence of an oxygen scavenger system (OSS) consisting of 1 mM Trolox (Fisher Scientific, Cat. No. AC218940050), 5 mM 3,4-Dihydroxybenzoic acid (Fisher Scientific, Cat. No. AC114891000), 50 nM protocatechuate dioxygenase (Millipore Sigma, Cat. No. P8279-25UN), and each sample well was imaged by objective-TIRF microscopy.

### Single-molecule fluorescence microscopy

Initial optimizations of auxiliary probe design, FP sequences, and imaging temperature (**Fig. 2**, **Supplementary Fig. 2b,c**) as well as quantification experiments in genomic DNA (**Fig. 5**) were performed using an Olympus IX-81 objective-type TIRF (O-TIRF) microscope with a 60× oil-immersion objective (APON 60XOTIRF, 1.45 NA) equipped with both a cellTIRF^TM^ illuminator and a z-drift control module (Olympus IX2-ZDC2). An EMCCD (electron-multiplying charge-coupled device) camera (Andor IXon 897, EM gain 150) was used to record the movies. For FRET measurements, Cy3 was excited by a 532 nm laser (OBIS 637 nm LX, 100 mW at a power of 9 mW as measured at the objective) and a calculated evanescent field penetration depth of 80 nm, and Cy5 fluorescence detected after passing through a dichroic mirror (ZT405/488/532/640rpc, Chroma) and an emission filter (ET705/100m, Chroma). The signal integration time (exposure time) per frame was 500 ms unless otherwise noted, and movies 2-10 min in length (in the final assay, 2 min) were collected per FOV. An objective heater (BIOPTECHS) was used to control the imaging temperature (at 26.5 °C, in the final assay) after calibration with an infrared thermometer (Lasergrip 800, Etekcity).

For further optimizations of FP concentration and CP design as well as initial calibration curves (**Fig. 3**, **Fig. 4**, **Supplementary Fig. 2**, and **Supplementary Fig. 3**), we used an Olympus IX-81 O-TIRF microscope equipped with a cellTIRF^TM^ illuminator, an ASI CRISP Z-drift control modules, and an EMCCD camera (Evolve 512, Photometrics). Cy3 was excited by TIRF with a 532 nm laser at a power of 9 mW, and Cy5 emission detected after passing through a Cy3-A647 FRET dichroic mirror (ZT40DRC-UF2, Chroma) and an emission filter (ET655LP-TRF filter, Chroma). An objective heater (BIOPTECHS) was used to control the temperature following the same calibration procedure.

Consecutive multiple-FOV imaging was performed using a journal programmed in Metamorph^39^ (**Supplementary Fig. 5**). A total of 10 FOVs were collected for all quantification experiments (**Fig. 4** and **Fig. 5**).

### Processing and analysis of objective-TIRF data

A set of custom MATLAB codes were used to identify spots with significant intensity fluctuations within each FOV, generate intensity-versus-time traces at each spot, fit these traces with a two-state hidden Markov model (HMM) to generate idealized traces, and eventually identify and characterize transitions with idealized traces. A set of filtering criteria were automatically generated to distinguish target-specific signal from non-specific background signal by feeding traces from no-target control experiments and unmethylated target-only experiments as negative training datasets, and traces from methylated target-only experiments as positive training datasets, into a SiMREPS optimizer (see **Supplementary Note 4** and **Supplementary Table 2**). Detailed discussions of the data analysis pipeline are published elsewhere^35, 36, 39, 49, 50^.

### Compilation of blood DNA methylation using recountmethylation

Recountmethylation is a R/Bioconductor package with 12, 537 uniformly processed EPIC and HM450K blood samples from GEO^42, 51^. All data on GEO as of December of 2022 measured by Illumina EPIC array and Illumina HM450K array are compiled and made available on its public data server (https://recount.bio/data/). To extract all DNA methylation beta values for the 102 nt BCAT1 promoter measured by Illumina EPIC array, a custom-written R code was used to extract and analyze data from the recountmethylation data server (see https://github.com/dai905/My_Recountmethylation/blob/ed415fac70776f33e1d57e098989a278f8e216f1/B lood_EPIC_v1.Rmd). Compiled data is available on the online server (see https://recount.bio/data/remethdb_h5se-gm_epic_0-0-2_1589820348/ and https://recount.bio/data/remethdb_epic-hm850k_h5se_gm_1669220613_0-0-3/).

### Compilation of DNA methylation in blood cells from UCSC genome browser

Initially, a total of 121 blood-related tracks were selected by manually screening UCSC Track Hubs using the search term “DNA methylation” for genome assembly hg38 (see publicly available UCSC genome browser session: https://genome.ucsc.edu/s/dai905/hg38_bloodcell_121). A custom-written R code was used to filter and combine methylation levels available at BCAT1 promoter. Eventually, a total of 72 tracks across various types of blood cells were selected.

### Targeted bisulfite pyrosequencing

Genomic DNA was transferred to the UM Epigenomics Core for pyrosequencing assay. The samples were quantitated using the Qubit BR dsDNA kit (ThermoFisher, Cat. No. Q32850). A total of 500 ng of genomic DNA was used for bisulfite conversion using the Zymo EZ DNA Methylation kit (Zymo Research, Cat. No. D5001), according to the manufacturer’s instructions. The converted DNA was eluted at a 10 ng/µL concentration for PCR amplification of region of interest. PCR amplification was done with Qiagen’s HotStarTaq Master Mix Kit (Qiagen, Cat. No. 203443), with 20 ng of converted DNA and 0.2 µM for each of primers in 30 µL final volume. The primers used for PCR amplification were GGTTGGGAGAGATTTTATTATTTGG (forward) and ATCCCCACTACAACAAAACCTAAA (reverse, biotinylated), using the following program: (i) heat activation at 95 °C for 15 min; (ii) amplification with 35 cycles of denaturation at 94 °C for 30 s, annealing at 50 °C for 30 s and elongation at 72 °C for 1 min; (iii) Final extension at 72 °C for 1 min; (iv) holding at 4 °C. PCR products size was verified on the Tapestation 4200 HS D1000 screentape assay. Samples were then processed for pyrosequencing on the Pyromark Q96 ID according to the manufacturer’s instructions. The sequencing primer used was GGTTTGGGGGAGTAG.

## Acknowledgments

We thank Dr. Damon Hoff of the Single Molecule Analysis in Real-Time (SMART) Center at the University of Michigan for assistance and advice with instrumentation, Dr. Shankar Mandal, now at the Thermo Fisher, for initial drafting, and Dr. Claudia Lalancette of the Biomedical Research Core Facilities (BRCF) ’s Epigenomics Core at the University of Michigan for designing and completing the targeted bisulfite pyrosequencing assay. This work was supported by NIH grant R21 CA225493 (N.G.W. and M.T.).

## Author contributions

L.D. and A.J.-B. conceived and designed the study. L.D. performed all experiments and analyzed all the data except the targeted bisulfite pyrosequencing. L.D. wrote the manuscript with input from all authors. A.J.-B. contributed to major revision of the manuscript. N.G.W. supervised the entire study. M.T. and P.W.L. and A.J.-B. supervised parts of the study.

## Inclusion and diversity

We support inclusive, diverse, and equitable conduct of research.

