## Supplementary Information for "Ultrasensitive amplification-free quantification of a methyl CpG-rich cancer biomarker by single-molecule kinetic fingerprinting"

The manuscript was written based on contributions from all authors.

All authors have given approval to the final version of the manuscript.

#### Supplementary Note 1: Optimization of imaging temperature, FP concentration and capture probes

Due to the low background signal, our sensitivity is primarily limited by the number of molecules that can be detected within the measurement time. Therefore, to achieve higher sensitivity, we imaged 10 FOVs per sample well and used the sum of all accepted counts across the 10 FOVs as our readout for that sample well (**Supplementary Fig. 5**). To ensure that this sequential multi-FOV acquisition does not extend measurement time more than necessary, we optimized imaging temperature and FP concentration to maximize the expected number of binding and dissociation transitions ($N_{b+d}$) per unit time, using the 102 nt MBC Mimic and capture probes CP1 + CP2 (**Supplementary Fig. 2a**). We first optimized imaging temperature from room temperature (RT, ~22 °C) up to 28 °C (**Supplementary Fig. 2b,c**), reasoning that elevated temperature should accelerate the kinetics of many reactions, resulting in a higher $N_{b+d}$ due to a lower $\tau_{on}$ and possibly lower $\tau_{off}$. From RT to 26.5 °C, $N_{b+d}$ indeed increased due to a shorter $\tau_{on}$, indicating destabilization of interactions between FPs and their docking sites (**Supplementary Fig. 2b,c**). However, above 26.5 °C, $N_{b+d}$ started to decline due to a longer $\tau_{off}$ (**Supplementary Fig. 2b,c**); this could be explained by a reduced rate of FP binding at higher temperatures, an increasing proportion of binding events too short to detect at our time resolution, or partial dissociation of auxiliary probes at high temperatures (since the $T_m$ of Aux1 is only 33 °C), placing even bound pairs of FPs out of FRET distance. In addition to imaging temperature, we optimized the concentration of the two FPs, which is expected to be directly proportional to the apparent (pseudo-first order) rate constant of binding, which is the reciprocal of $\tau_{off}$, resulting in higher $N_{b+d}$. Indeed, our experiments (**Supplementary Fig. 2d**) suggested an upshift in $N_{b+d}$ distribution as FP concentration increased. However, we decided not to use concentrations higher than 100 nM because S/N declined significantly above this concentration (**Supplementary Fig. 2e**). In order to reach the same sensitivity as that seen at 100 nM FPs, the S/N threshold of our analysis would have had to be reduced, resulting in more false positives being accepted in no-target control experiments (**Supplementary Fig. 7**), which would have defeated the purpose of increasing sensitivity.

Apart from imaging conditions, sensitivity also depends on capture efficiency which is a function of capture probe design and capture protocol. In particular, we reasoned that a second capture probe might increase capture efficiency (and sensitivity) by accelerating capture kinetics and ensuring that even targets with one capture-binding sequence missing or damaged (e.g., by bisulfite treatment) could still be captured. Accordingly, we designed two capture probes: CP1, which binds the target at its 5′-end, and CP2, which binds the target at its 3′-end (**Supplementary Fig. 2a**). This double capture-probe scheme was utilized for all optimizations of imaging conditions (**Supplementary Fig. 2**). However, further quantification experiments suggested only modest improvements to sensitivity and reduced specificity with double capture (**Supplementary Fig. 3**), and the fact that CP1 occasionally generated inconsistent counts due to apparent degradation under long-term storage (**Supplementary Fig. 3d,e**) became a technical concern. Due to these concerns with CP1, we decided to use CP2 as the sole capture probe in our final construct as well as in the following characterization experiments.

#### Supplementary Note 2: Calculation of theoretical and apparent discrimination factors and specificity

The free energy of hybridization of all DNA complexes under final assay conditions were predicted using NUPACK (http://www.nupack.org/partition/new) and the conditions indicated below:

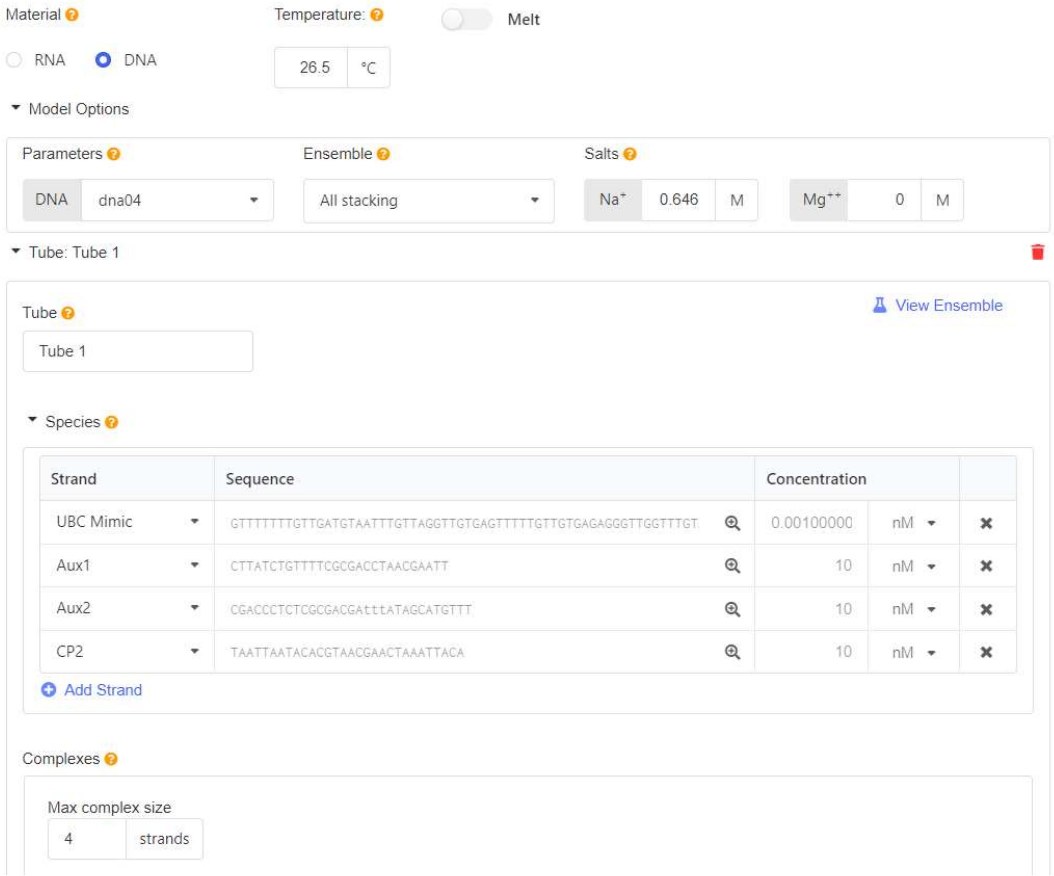


Since GU mismatches are not explicitly handled in the thermodynamic model, we treated them as GT mismatches and substituted them accordingly as shown in the "Sequence" cell above.

For similar reasons, we did not consider the effects of methyl groups on binding free energies, which have a much smaller influence on hybridization thermodynamics than nucleobase identity. All strand concentrations were set as shown above, temperature was set to 26.5 °C, and [$Na^+$] was set to 0.646 M to approximate the total concentration of monovalent cations in 4× PBS.

The theoretical maximum discrimination factor permitted by thermodynamics can be calculated as below[1]:

$$Q_{max,therm} = e^{-\Delta\Delta G^{\circ}/RT}$$

where

$$\Delta\Delta G^{\circ} = \Delta G^{\circ}_{MBC\ or\ MBC\ Mimic+Aux1+Aux2+C} - \Delta G^{\circ}_{UBC\ or\ UBC\ Mimic+Aux1+Aux2+CP2}$$

The apparent discrimination factor and specificity are defined as below:

$$Q_{app} = Discrimination\ factor = \frac{Total\ Negatives}{False\ Positives} = \frac{1}{1 - Specificity}$$

$$Specificity = \frac{True\ Negatives}{True\ Negatives + False\ Positives}$$

In a given experiment, the False Positives term is simply the number of accepted counts detected in the presence of only UBC/UBC Mimic at a certain concentration *C*. True Positives are equal to the total accepted counts detected in a mixture of MBC + UBC or MBC Mimic + UBC Mimic minus the False Positives detected in the presence of only UBC/UBC Mimic at concentration *C*.

Since our sensor is not designed to detect UBC/UBC Mimic, we define Total Negatives as the number of counts that would result if UBC/UBC Mimic were detected with the same sensitivity as MBC/MBC Mimic, proportional to the relative concentrations of unmethylated to methylated target. Therefore, the Total Negatives term is estimated as:

$$Total\ Negatives = Molar\ ratio * True\ Positives$$

where molar ratio equals $C_{UBC}/C_{MBC}$ or $C_{UBC\ Mimic}/C_{MBC\ Mimic}$. Finally,

$$Specificity = \frac{True\ Negatives}{True\ Negatives + False\ Positives} = \frac{True\ Negatives}{Total\ Negatives}$$

$$= \frac{Total\ Negatives - False\ Positives}{Total\ Negatives}$$

$$= 1 - \frac{False\ Positives}{Total\ Negatives}$$

$$= 1 - \frac{False\ Positives}{Molar\ ratio * True\ Positives}$$

$$= 1 - \frac{False\ Positives}{Molar\ ratio * (Total\ accepted\ counts - False\ Positives)}$$

#### Supplementary Note 3: Calculation of $T_m$ (melting temperature)

The $T_m$ values of oligonucleotide hybridization reactions were calculated using NUPACK according to settings shown in the screenshot below:

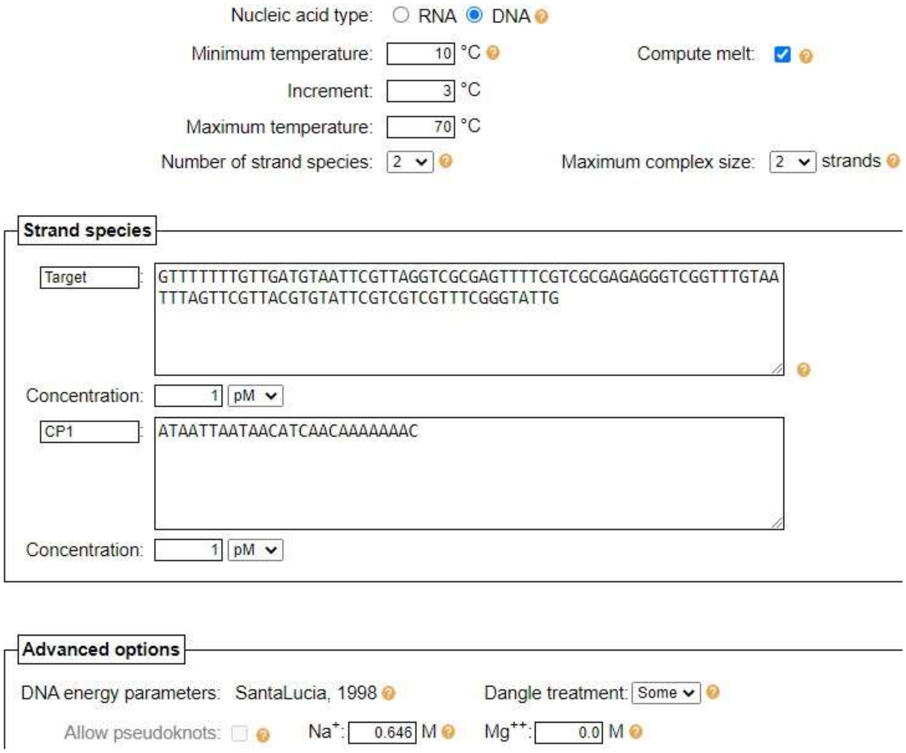


The target sequence shown here represents the 102 nt MBC Mimic where, as discussed in **Supplementary Note 2**, all uracils are replaced by thymines due to input restrictions. Calculations of the $T_m$ of CP binding were performed assuming a CP concentration of 1 pM, and calculations of the $T_m$ of FP binding were performed assuming a concentration of 100 nM for each FP. Each simulation generated a melting curve along with predicted concentrations of different complexes at specific temperatures. Simulated melting curves of different capture probes and auxiliary probes to the Target are shown in **Supplementary Fig. 10**.

#### Supplementary Note 4: Optimization of kinetic filtering criteria

Details regarding data analysis, including the SiMREPS data processing pipeline, are discussed at length in our previously published papers[2–4]. Briefly, two MATLAB programs called SiMREPS analysis suite and SiMREPS optimizer (http://inventions.umich.edu/technologies/6250-1_simreps-analysis-software-v-1-0, along with a user guide and links to example input and output files)[4] are used in BSM-SiMREPS. SiMREPS analysis suite consists of (1) a spot finder that recognizes single-molecule spots within a FOV and generates intensity-time traces from them according to a set of trace generation parameters, and (2) an analyzer that applies a set of kinetic filtering criteria (KFC) that can either be manually customized or automatically generated by a SiMREPS optimizer to maximize true positive counts and minimize false positive counts. Each experiment comes with a set of KFC and the optimized KFC is shown in **Supplementary Table 2** and used for all quantification experiments.

**Supplementary Table 1.** DNA oligonucleotide names, sequences, and descriptions. All sequences are listed 5′-to-3′.

| Name | Sequences | Description |
|---|---|---|
| BCAT1 Forward | GTCTTCCTGCTGATGCAATCCGCTAGGTCGCGAGTCTCCGCCGCGAGAGGGCCGGTCTGCAATCCAGCCCGCCACGTGTACTCGCCGCCGCCTCGGGCACTG | Full-length BCAT1 promoter forward strand, directly purchased from IDT |
| BCAT1 Reverse | CAGTGCCCGAGGCGGCGGCGAGTACACGTGGCGGGCTGGATTGCAGACCGGCCCTCTCGCGGCGGAGACTCGCGACCTAGCGGATTGCATCAGCAGGAAGAC | Full-length BCAT1 promoter reverse strand, directly purchased from IDT |
| dsBCAT1 | (Hybrid of BCAT1 Forward + BCAT1 Reverse) | ~1 μM dsBCAT1 was prepared by mixing equal amount of BCAT1 Forward and Reverse in a final buffer of 2× PBS, followed by heating at 90 °C for 5 min, annealing at 37 °C for 5 min, cooling down at room temperature. 10 μL aliquots were made and stored at -20 °C. |
| Me-BCAT1 Forward | GTCTTCCTGCTGATGCAATC/iMe-dC/GCTAGGT/iMe-dC/G/iMe-dC/GAGTCTC/iMe-dC/GC/iMe-dC/G/iMe-dC/GAGAGGGC/iMe-dC/GGTCTGCAATCCAGCC/iMe-dC/GCCA/iMe-dC/GTGTACT/iMe-dC/GC/iMe-dC/GC/iMe-dC/GCCT/iMe-dC/GGGCACTG | Full-length methylated BCAT1 promoter forward strand, directly purchased from IDT |
| Me-BCAT1 Reverse | CAGTGCC/iMe-dC/GAGG/iMe-dC/GG/iMe-dC/GG/iMe-dC/GAGTACA/iMe-dC/GTGG/iMe-dC/GGGCTGGATTGCAGAC/iMe-dC/GGCCCTCT/iMe-dC/G/iMe-dC/GG/iMe-dC/GGAGACT/iMe-dC/G/iMe-dC/GACCTAG/iMe-dC/GGATTGCATCAGCAGGAAGAC | Full-length methylated BCAT1 promoter reverse strand, directly purchased from IDT |
| dsMe-BCAT1 | (Hybrid of Me-BCAT1 Forward + Me-BCAT1 Reverse) | ~1 μM dsMe-BCAT1 was prepared by mixing equal amount of Me-BCAT1 Forward and Reverse in a final buffer of 2× PBS, followed by heating at 90 °C for 5 min, annealing at 37 °C for 5 min, cooling down at room temperature. 10 μL aliquots were made and stored at -20 °C. |

| | | |
|---|---|---|
| 10Me-BCAT1 Forward | GTCTTCCTGCTGATGCAATCCGCTAGGTCGCGAGTCTC/iMe-dC/GC/iMe-dC/G/iMe-dC/GAGAGGGC/iMe-dC/GGTCTGCAATCCAGCC/iMe-dC/GCCA/iMe-dC/GTGTACT/iMe-dC/GC/iMe-dC/GC/iMe-dC/GCCT/iMe-dC/GGGCACTG | Full-length methylated BCAT1 promoter (omitting methylation of the first three CpGs) forward strand, directly purchased from IDT |
| | **Non-mimic targets** | |
| 102 nt MBC | GTUTTUUTGUTGATGUAATU/iMe-dC/GUTAGGT/iMe-dC/G/iMe-dC/GAGTUTU/iMe-dC/GU/iMe-dC/G/iMe-dC/GAGAGGGU/iMe-dC/GGTUTGUAATUUAGUU/iMe-dC/GUUA/iMe-dC/GTGTAUT/iMe-dC/GU/iMe-dC/GU/iMe-dC/GUUT/iMe-dC/GGGUAUTG | The expected forward strand sequence of product after treating dsMe-BCAT1 with Methylation-Lightning$^{TM}$ Kit (in principle only the forward sequence will be captured, so the reverse strand is not shown here, but is identical to 102 nt rMBC) |
| 102 nt UBC | GTUTTUUTGUTGATGUAATUUGUTAGGTUGUGAGTUTUUGUUGUGAGAGGGUUGGTUTGUAATUUAGUUUGUUAUGTGTAUTUGUUGUUGUUTUGGGUAUTG | The forward strand sequence of product after treating dsBCAT1 with Methylation-Lightning$^{TM}$ Kit |
| 102 nt fMBC | GTUTTUUTGUTGATGUAATU/iMe-dC/GUTAGGT/iMe-dC/G/iMe-dC/GAGTUTU/iMe-dC/GU/iMe-dC/G/iMe-dC/GAGAGGGU/iMe-dC/GGTUTGUAATUUAGUU/iMe-dC/GUUA/iMe-dC/GTGTAUT/iMe-dC/GU/iMe-dC/GU/iMe-dC/GUUT/iMe-dC/GGGUAUTG | Sequence of product after treating Me-BCAT1 Forward with Methylation-Lightning$^{TM}$ Kit. Identical to the forward strand of 102 nt MBC |
| 102 nt rMBC | UAGTGUU/iMe-dC/GAGG/iMe-dC/GG/iMe-dC/GG/iMe-dC/GAGTAUA/iMe-dC/GTGG/iMe-dC/GGGUTGGATTGUAGAU/iMe-dC/GGUUTUT/iMe-dC/G/iMe-dC/GG/iMe-dC/GGAGAUT/iMe-dC/G/iMe-dC/GAUUTAG/iMe-dC/GGATTGUATUAGUAGGAAGAU | Sequence of product after treating Me-BCAT1 Reverse with Methylation-Lightning$^{TM}$ Kit |

| | | |
|---|---|---|
| 10Me-fMBC | GTUTTUUTGUTGATGUAATUUGUTAGGTUGUGAGTUTU/iMe-dC/GU/iMe-dC/G/iMe-dC/GAGAGGGU/iMe-dC/GGTUTGUAATUUAGUU/iMe-dC/GUUA/iMe-dC/GTGTAUT/iMe-dC/GU/iMe-dC/GU/iMe-dC/GUUT/iMe-dC/GGGUAUTG | Bisulfite conversion product of 10Me-BCAT1 Forward |
| | **Mimic targets** | |
| 102 nt MBC Mimic | GTUTTUUTGUTGATGUAATUCGUTAGGTCGCGAGTUTUCGUCGCGAGAGGGUCGGTUTGUAATUUAGUUCGUUACGTGTAUTCGUCGUCGUUTCGGGUAUTG | The mimic for 102 nt MBC without methyl modifications, directly purchased from IDT |
| 102 nt UBC Mimic | GTUTTUUTGUTGATGUAATUUGUTAGGTUGUGAGTUTUUGUUGUGAGAGGGUUGGTUTGUAATUUAGUUUGUUAUGTGTAUTUGUUGUUGUUTUGGGUAUTG | The mimic for 102 nt UBC, directly purchased from IDT |
| 55 nt MBC Mimic | GTUTTUUTGUTGATGUAATUCGUTAGGTCGCGAGTUTUCGUCGCGAGAGGGUCGG | The mimic for 55 nt MBC without methyl modifications, directly purchased from IDT |
| 55 nt UBC Mimic | GTUTTUUTGUTGATGUAATUUGUTAGGTUGUGAGTUTUUGUUGUGAGAGGGUUGG | The mimic for 55 nt UBC, directly purchased from IDT |
| 42 nt MBC Mimic | TGUAATUCGUTAGGTCGCGAGTUTUCGUCGCGAGAGGGUCGG | The mimic for 42 nt MBC without methyl modifications, directly purchased from IDT |
| 42 nt UBC Mimic | TGUAATUUGUTAGGTUGUGAGTUTUUGUUGUGAGAGGGUUGG | The mimic for 42 nt UBC, directly purchased from IDT |
| | **Sensor construct** | |
| Aux1 | **CTTATCTGT**TTTCGCGACCTAACGAATT | Auxiliary probe 1, providing docking site for FP1, 1b and 1c |
| Aux2 | CGACCCTCTCGCGACGATTT**ATAGCATG**TTT | Auxiliary probe 2, providing docking site for FP2 and 2c |
| Biotin-Aux2 | CGACCCTCTCGCGACGATTT**ATAGCATG**TTT/3bioTEG/ | Biotinylated auxiliary probe 1, providing docking site for FP2 and also serving as a capture probe in Fig. 2. |
| CP1 | /5Biosg/ATAATTAATAACATCAACAAAAAAAC | Capture probe 1, used in Fig. 2 and Fig. 3. |

| | | |
|---|---|---|
| CP1-LNA | /5Biosg/ATAATTAATAA+CAT+CAA+CAA+AAA+AA+C | Locked nucleic acid(LNA)-modified CP1, used in Supplementary Fig. 3. |
| CP2 | /5Biosg/TAATTAATACACGTAACGAACTAAATTACA | Capture probe 2, used in Fig. 3, 4 and 5. |
| Block1 | TCACAACCTAACAAATTACA | Blocker 1, a short strand complementary to UBC or UBC Mimic (all sizes) at the Aux1-binding region. |
| Block2 | CAACCCTCTCACAACAA | Blocker 2, a short strand complementary to UBC or UBC Mimic (all sizes) at the Aux2-binding region. |
| | **Fluorescent Probes (FPs)** | |
| FP1 | /5Cy3/CAGATAAG | Fluorescent probe 1, interacting with Aux1 *via* 8-nt binding region |
| FP1b | /5Cy3/ACAGATAAG | Fluorescent probe 1b, interacting with Aux1 *via* 9-nt binding region |
| FP1c | /5Cy3/TTACAGATAAG | Fluorescent probe 1c, interacting with Aux1 *via* 9-nt binding region |
| FP2 | CATGCTAT/3Cy5Sp/ | Fluorescent probe 2, interacting with Aux2 *via* 8-nt binding region |
| FP2c | CATGCTATTTT/3Cy5Sp/ | Fluorescent probe 2c, interacting with Aux1 *via* 8-nt binding region and 3T linker |
| | **Genomic DNA** | |
| WGA | NA | (Directly purchased from Zymo Research, cat. no. D5013-1) Human HCT116 DKO Non-methylated DNA, generated using phi29 DNA polymerase based whole genome amplification techniques from HCT116 DKO cell line derived genomic DNA |
| +Me WGA | NA | (Directly purchased from Zymo Research, cat. no. D5013-2) Human WGA Methylated DNA, generated using human WGA Non-methylated DNA that has |

| | | |
|---|---|---|
| | | been enzymatically methylated at all double-stranded CG dinucleotides using M.SssI methyltransferase. |
| Blood DNA | NA | (Directly purchased from Enzo Lifesciences, ENZ-GEN117-0100) Human Genomic DNA, Male, comprised of a pool of ≥ 8 normal donors. It is intact DNA extracted from freshly harvested whole blood of healthy males. DNA is treated with DNase-free RNase to remove residual contaminant RNA. |
| BS WGA | NA | Bisulfite conversion product of WGA |
| BS +Me WGA | NA | Bisulfite conversion product of +Me WGA |
| BS Blood DNA | NA | Bisulfite conversion product of Blood DNA |
| BS Epi | NA | (Directly purchased from Qiagen, cat. no. 59695) Epitect bisulfite converted unmethylated PCR Control DNA |
| BS +Me Epi | NA | (Directly purchased from Qiagen, cat. no. 59695) Epitect bisulfite converted methylated PCR Control DNA |
| **Bisulfite Pyrosequencing** | | |
| BCAT1_Pyro_FWD | GGTTGGGAGAGATTTTATTATTTGG | Forward primer for PCR amplification of bisulfite-converted BCAT1 promoter in genomic DNAs, purchased from IDT |
| BCAT1_Pyro_BioREV | /5Biosg/ATCCC**C**ACTACAACAAAACCTAAA | Biotinylated reverse primer for PCR amplification of bisulfite-converted BCAT1 promoter in genomic DNAs, purchased from IDT; a single mismatch in highlighted in red |
| BCAT1_Pyro_Seq | GGTTTGGGGGAGTAG | Sequencing primer for PCR-amplified bisulfite-converted BCAT1 promoter in genomic DNAs used for pyrosequencing, purchased from IDT |

**Supplementary Table 2.** Optimized parameter sets for trace generation and analysis.

| Trace Generation Parameters | |
|---|---|
| use fluctuation map? | 2 ('Nb+d map') |
| Stdfactor | 15 |
| start frame | 1 |
| end frame | 240 |
| edgePx | 20 |
| Percentilecut | 0.95 |
| ROI size (pixels) | 3 |

| Trace Analysis Parameters (kinetic filtering criteria, KFC) | |
|---|---|
| start frame | 1 |
| end frame | 240 |
| exposure time (s) | 0.5 |
| Smoothframes | 1 |
| remove_single_frame_events | FALSE |
| Ithresh | 151.6923572 |
| SNthresh | 2 |
| SNthresh_trace | 2.5 |
| min_Nbd | 12 |
| max_Nbd | 65 |
| min_tau_on_median (s) | 0.5 |
| min_tau_off_median (s) | 0.5 |
| max_tau_on_median (s) | 7 |
| max_tau_off_median (s) | 35 |
| max_tau_on_cv | 1.8 |
| max_tau_off_cv | Inf |
| max_tau_on_event (s) | 8 |
| max_tau_off_event (s) | 80 |
| max_I_low_state | 194000 |
| vary_I_vals | FALSE |
| num_intensity_states | 2 |
| ignore_post_bleaching | FALSE |
| bleaching_wait_time (s) | Inf |
| use_FRET_threshold | FALSE |
| FRET_threshold | 0 |

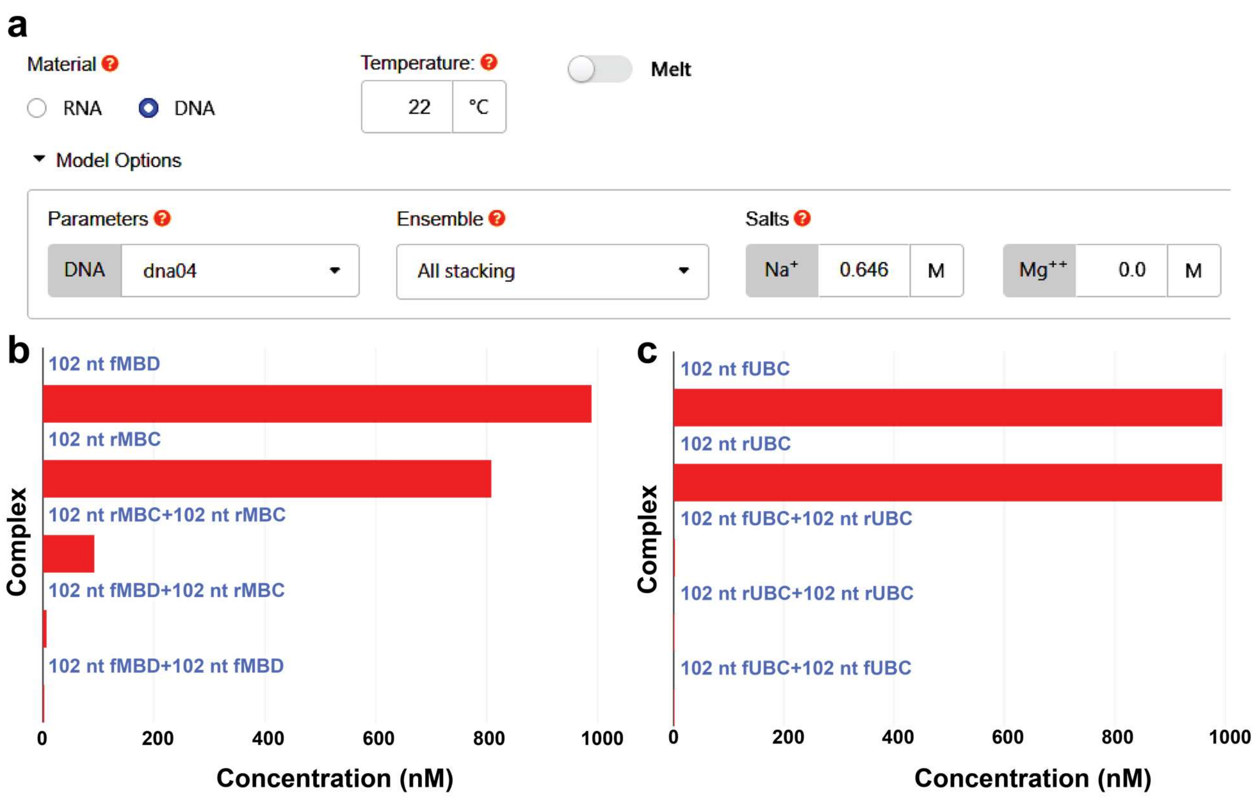


**Supplementary Fig. 1. | Nupack prediction of hybridization of Forward and Reverse strands after bisulfite conversion of methylated or unmethylated BCAT1.** See **Supplementary Note 2** for detailed description of prediction conditions. 102 nt fMBC and rMBC are sequences of bisulfite converted Me-BCAT1 Forward and Me-BCAT1 Reverse respectively; 102 nt fUBC and rUBC are sequences of bisulfite converted BCAT1 Forward and BCAT1 Reverse respectively. **a** Prediction condition for hybridization of Forward and Reverse strands after bisulfite conversion of methylated BCAT1. All input sequences with Us replaced by Ts are 1 μM. Maximum complex size is 2 strands. **b** Nupack prediction result using condition in panel a. **c** Prediction condition for hybridization of Forward and Reverse strands after bisulfite conversion of unmethylated BCAT1. **d** Nupack prediction result using condition in panel c.

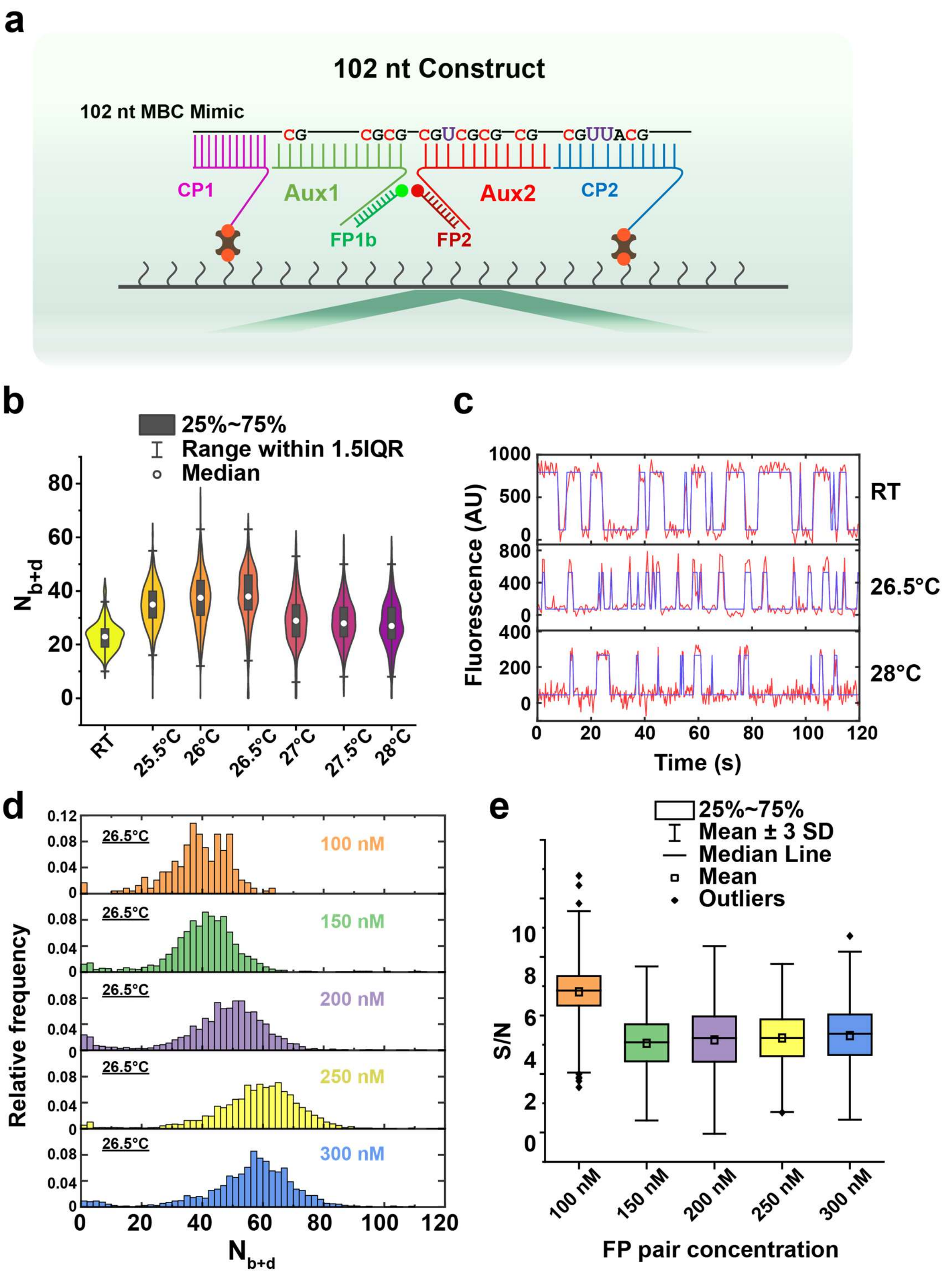


**Supplementary Fig. 2. | Optimization of imaging conditions**. **a** Assay construct used in optimizations. The 102 nt construct is designed for detecting 102 nt MBC Mimic with two biotinylated capture probes, CP1 and CP2. **b** Violin plots showing effect of imaging temperature on $N_{b+d}$ distribution. **c** Representative intensity-time traces (red lines) fit by HMM (blue lines) at different temperatures. **d** Histograms showing effects of FP concentration on $N_{b+d}$ distribution. **e** Effects of FP concentration on signal-to-noise ratio (S/N) using the same dataset as in panel d.

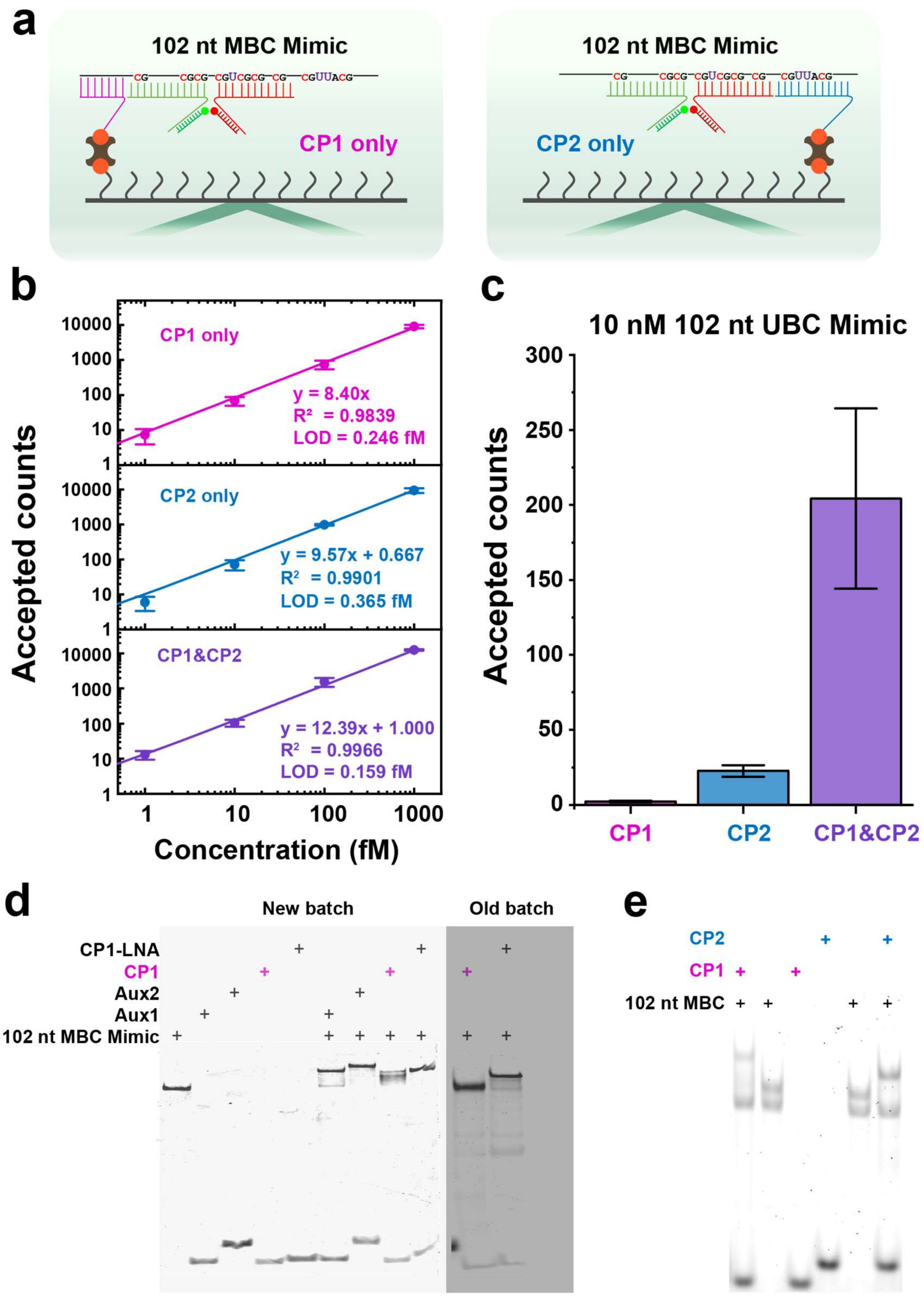


**Supplementary Fig. 3. | Analytical performance using different capture probe sets. a** Assay constructs using individual capture probes. **b** Standard curves of 102 nt MBC Mimic using different capture probe sets. **c** False positive counts using different capture probe sets. Data from panels b and c were collected in imaging conditions identical to those of Fig. 2. For each capture probe set, a separate set of kinetic filtering criteria were generated by our automated parameter optimizer so as to maximize true positive and minimize false positive counts. **d** 12% native PAGE showing evidence of degradation of CP1 and CP1-LNA under long-term storage. Each strand of new batch was loaded with the same amounts as of old batch. Comparing the two lanes for the old batch with the corresponding lanes of the new batch, the bands corresponding to free CP1 and CP1-LNA are much fainter in the old batch; furthermore, in the old batch there is no band shift indicative of CP1 binding to MBC Mimic, and a reduced shift for CP1-LNA binding to the MBC Mimic. **e** 5% native PAGE showing binding of CP1 and CP2 to 102 nt MBC. Both gels in Panel d-e are stained by SYBR Gold and visualized using the Cy2 fluorescence setting.

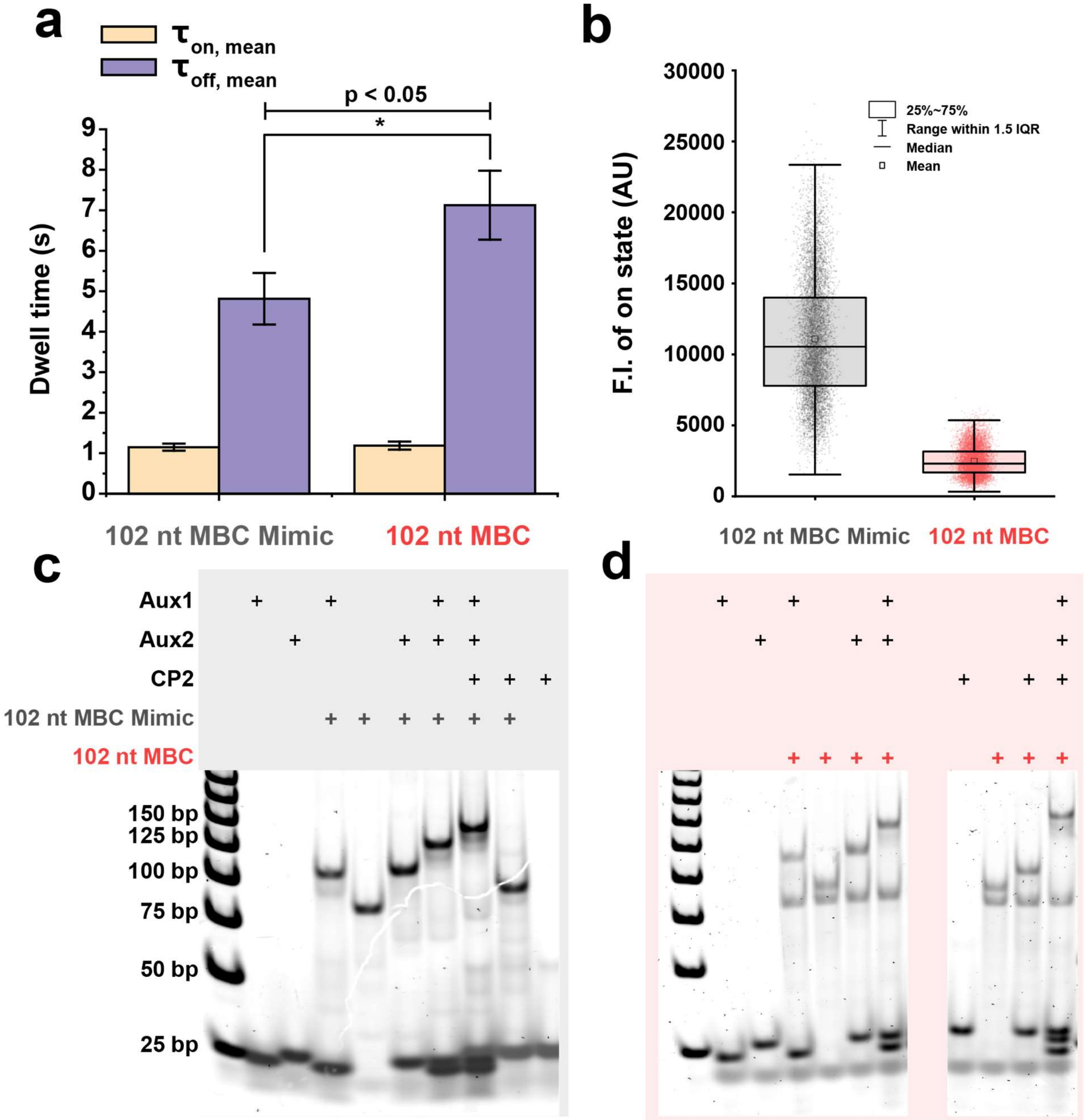


**Supplementary Fig. 4. | Differences between 102 nt MBC Mimic and 102 nt MBC.** See Fig. 3a for construct design and probe details. **a** Mean dwell times observed during detection of 102 nt MBC Mimic and 102 nt MBC. Mean $\tau_{on}$ and $\tau_{off}$ are calculated by fitting an exponential decay function to cumulative histograms of dwell times of individual events in all traces. Datapoints are presented as mean ± s.d. with n = 3 independent experiments. P-value as assessed using a single-tailed, unpaired two-sample t-test. **b** Distributions of bound-state intensity during detection of 102 nt MBC Mimic and 102 nt MBC. Experiments shown in panels a and b used the same capturing and imaging conditions as in Fig. 3. **c** 5% native PAGE showing the binding of probes to the 102 nt MBC Mimic. **d** 5% native PAGE to show probes' binding to 102 nt MBC. See **Supplementary Fig. 11** for the complete raw gel image of panel d.

### Multiple FOV data acquisition

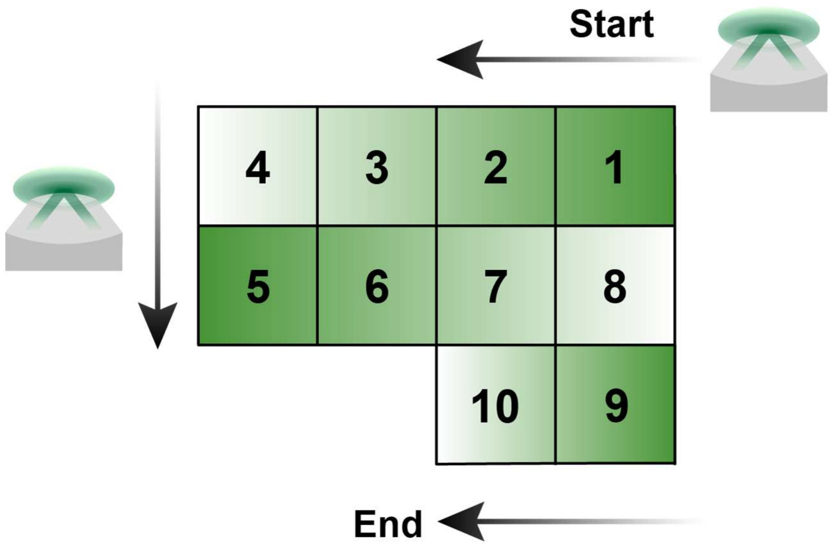


**Supplementary Fig. 5 | Acquisition scheme of multiple-FOV detection.** Serpentine-scan acquisition of 10 FOVs. Relative to the microscope stage, the objective moves from the top right to the left on the first row, moves down to the next row and then moves to the right, following the brightening color gradient shown in this panel. Eace square cell represents a single FOV.

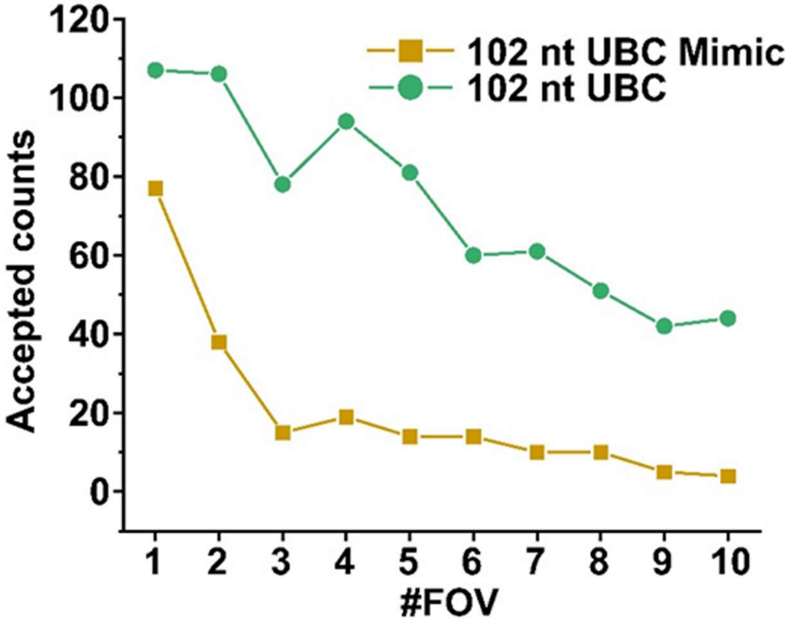


**Supplementary Fig. 6 | Accepted counts across multiple sequentially imaged FOVs in assays of samples containing 10 nM 102 nt UBC Mimic or 10 nM 102 nt UBC.** For both the mimic and the bisulfite-treated target, a clear decrease is observed in accepted counts over time, suggesting gradual dissociation of weakly bound construct containing UBC Mimic or UBC from the surface.

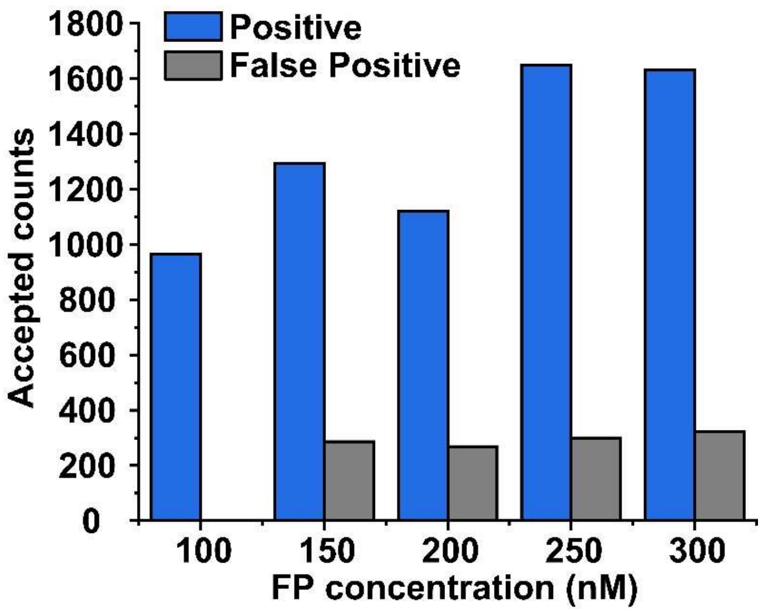


**Supplementary Fig. 7 | Specificity comparison among different concentrations of the FP pair FP1b + FP2.** Each value represents the concentration of *each* FP in the mixture; e.g., at 100 nM, there is 100 nM FP1b and 100 nM of FP2. At each concentration, a separate set of KFCs were generated by our automated parameter optimizer so as to maximize true positive and minimize false positive counts. Positive datasets comprise traces collected in the presence of 1 pM 102 nt MBC Mimic with varying concentrations of FP1b + FP2. The negative dataset comprises traces acquired in 100 nM FP1b and 100 nM of FP2 at 26.5 °C, in one FOV with each of three separate conditions: 1) no target control; 2) 1 nM 102 nt UBC Mimic; 3) 5 nM 102 nt UBC Mimic. The same negative dataset was used for training KFCs at varying concentrations of FP1b + FP2. Each datapoint was collected from a single FOV. False Positive represents total accepted counts of the entire negative dataset after applying optimized KFCs. Although a marginal increase in positive counts is observed across FP concentrations higher than 100 nM, it is accompanied by proportionally much larger increase in false positives: while nearly zero false positives were detected in 100 nM FPs, over 200 false positives were detected in just a single FOV (per negative condition) for each FP concentration $\geq$150 nM. In other words, any benefit from the marginal increase in signal amplitude is swamped by a significant decline in specificity.

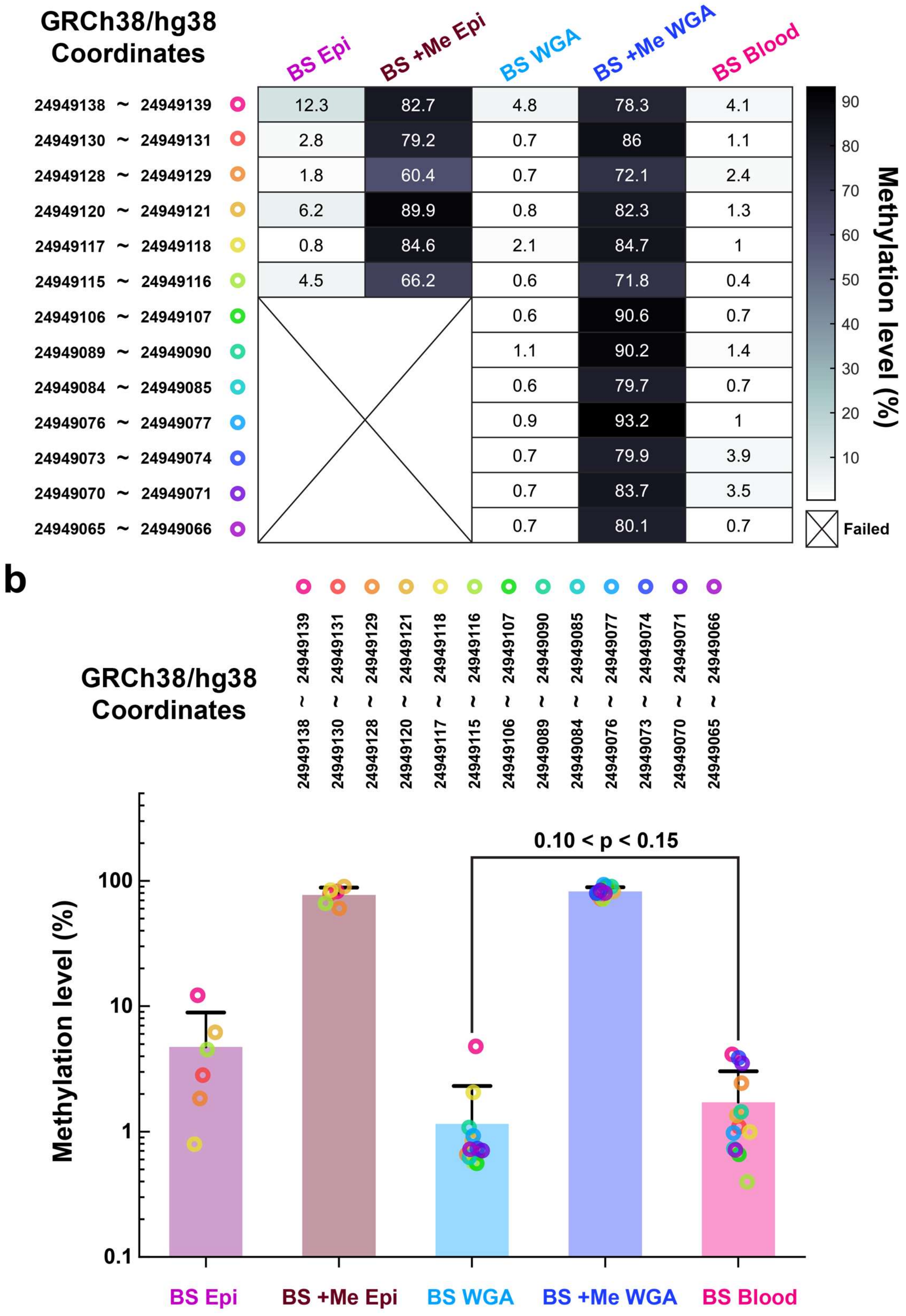


**Supplementary Fig. 8 | Results of targeted bisulfite pyrosequencing at 102 nt BCAT1 promoter across different genomic DNA samples.** CpG sites along with their coordinates are represented as circles with different colors. Genomic samples include: **BS Epi**, bisulfite-converted unmethylated DNA from Epitect; **BS +Me Epi**, bisulfite-converted methylated DNA from Epitect; **BS WGA**, bisulfite-converted whole-genome amplified DNA (see Fig. 5b); **BS +Me WGA**, bisulfite-converted methylated whole-genome amplified DNA (see Fig. 5b); and **BS Blood**, bisulfite-converted whole blood DNA, the same batch of sample as that analyzed in Fig. 5b. **a** Color-mapped visualization of methylation levels measured by targeted bisulfite pyrosequencing at 102 nt BCAT1 promoter. Three independent measurements were conducted for each type of sample; the value in each cell is the mean of any "passed" or "check" readouts excluding "failed" readouts. The status of each readout is determined by the built-in thresholds of pyrosequencer software. In BS Epi and BS +Me Epi, CpG sites for which all measurements failed are represented by a box with diagonal lines (see Methods for details.) **b** Distribution of methylation levels across different CpG sites in each type of sample, replotted from data in panel a. Each bar represents the mean value of available methylation readouts across different CpG sites. Error bars represent 1 standard deviation. P-value as assessed using a single-tailed, unpaired two-sample t-test.

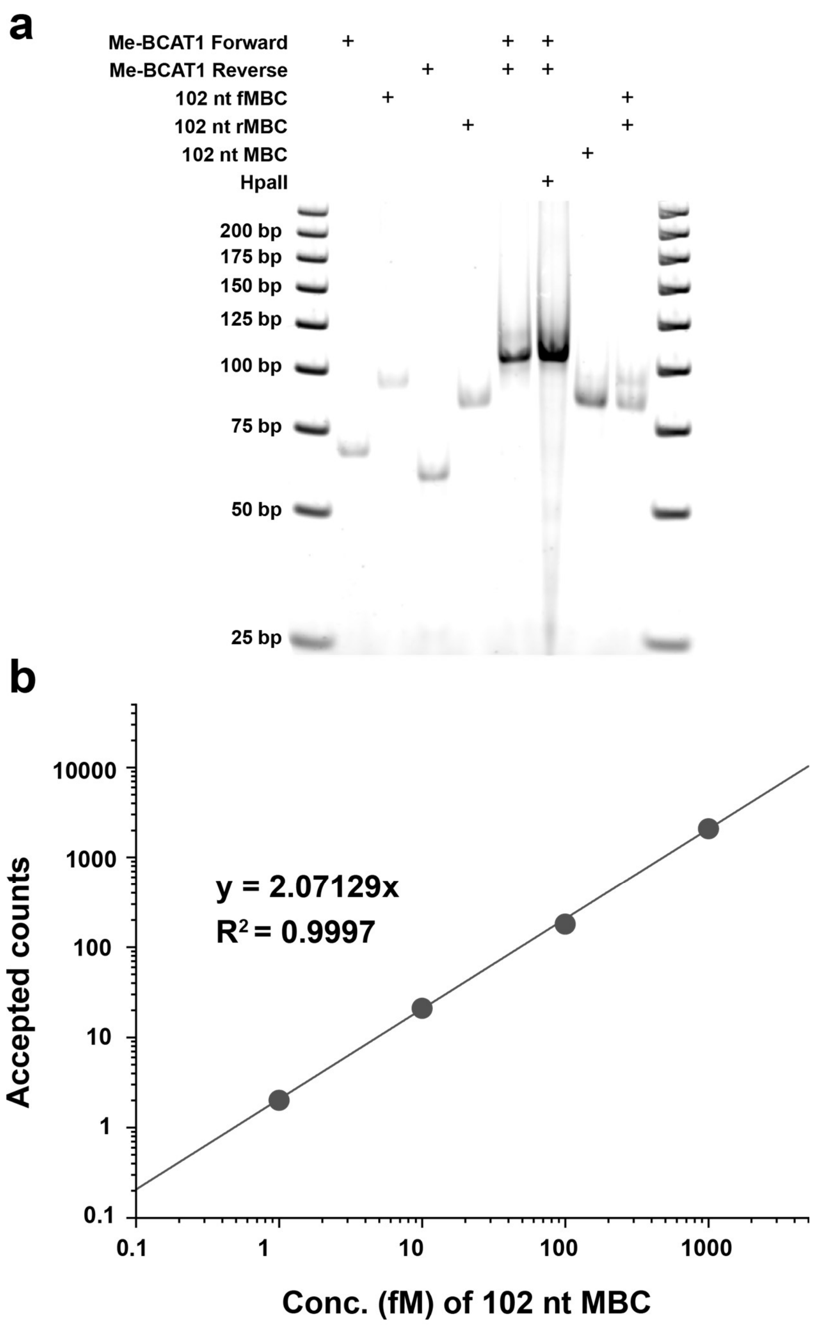


**Supplementary Fig. 9 | Evidence of biased yield of Forward and Reverse target strands in bisulfite conversion. a** 5% native PAGE to assess recovery of individual strands after bisulfite conversion (see **Supplementary Table 1** for details of each DNA strand). HpaII is a methylation-sensitive restriction enzyme whose digestion activity is inhibited by methylated CpGs. The third lane from the right (containing 102 nt MBC, the bisulfite conversion product of double-stranded 102 nt BCAT1) shows a much lower concentration of forward strand than reverse strand after bisulfite treatment and downstream column cleanup, despite equal inputs of the two strands into the reaction. **b** Quantification of different concentrations of 102 nt MBC using the same batch of samples shown in panel a. There is a significant (>4-fold) decrease in accepted counts compared to standard curves in Fig. 4. This is likely because the input concentration of Forward strand is significantly overestimated by Qubit since the majority of recovered ssDNA is the Reverse strand.

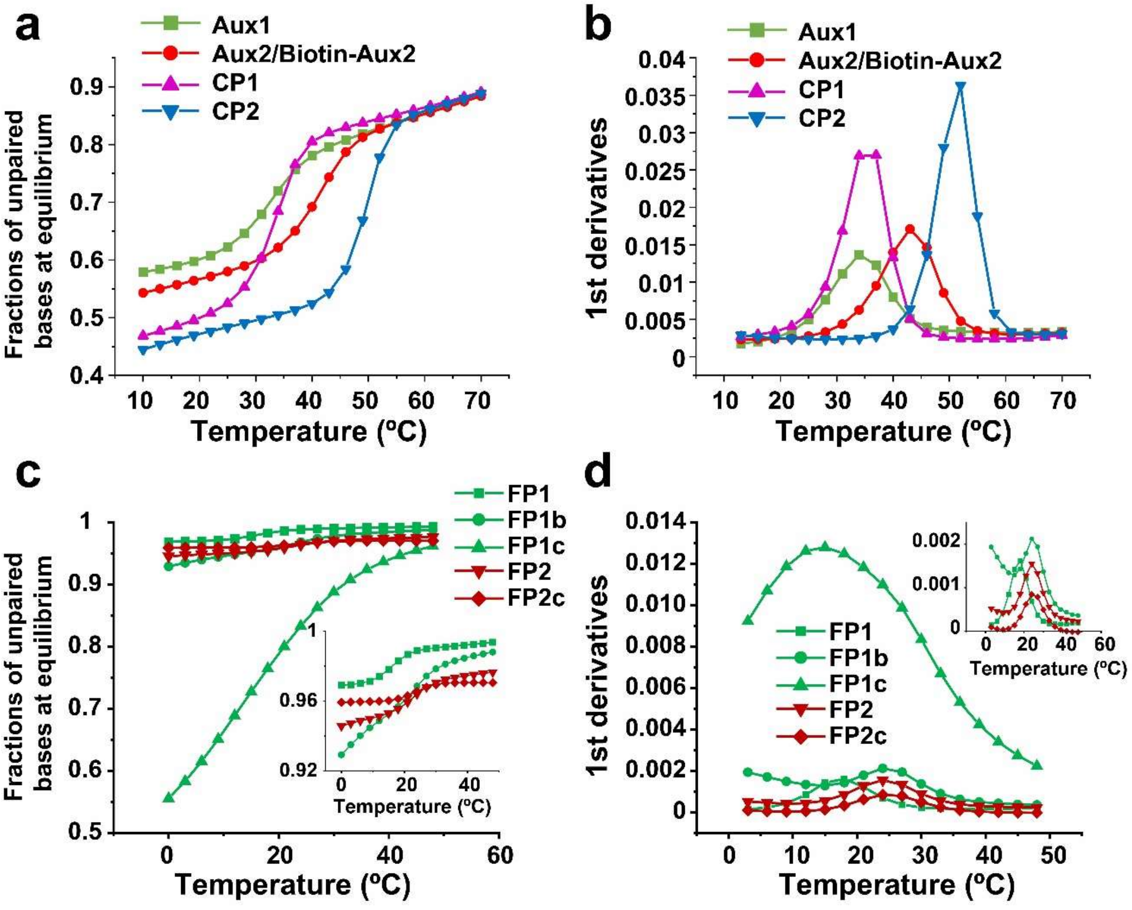


**Supplementary Fig. 10 | Predictions of $T_m$ by simulated melting curves. a,b** Melting curves (**a**) and first derivatives of unpaired base fraction (**b**) with respect to temperature for interactions of capture probes and auxiliary probes with 102 nt MBC Mimic. **c,d** Melting curves (**c**) and first derivatives (**d**) of unpaired base fraction with respect to temperature for interactions of different FPs with their respective auxiliary probes. Simulated melting curves were generated using NUPACK and replotted.

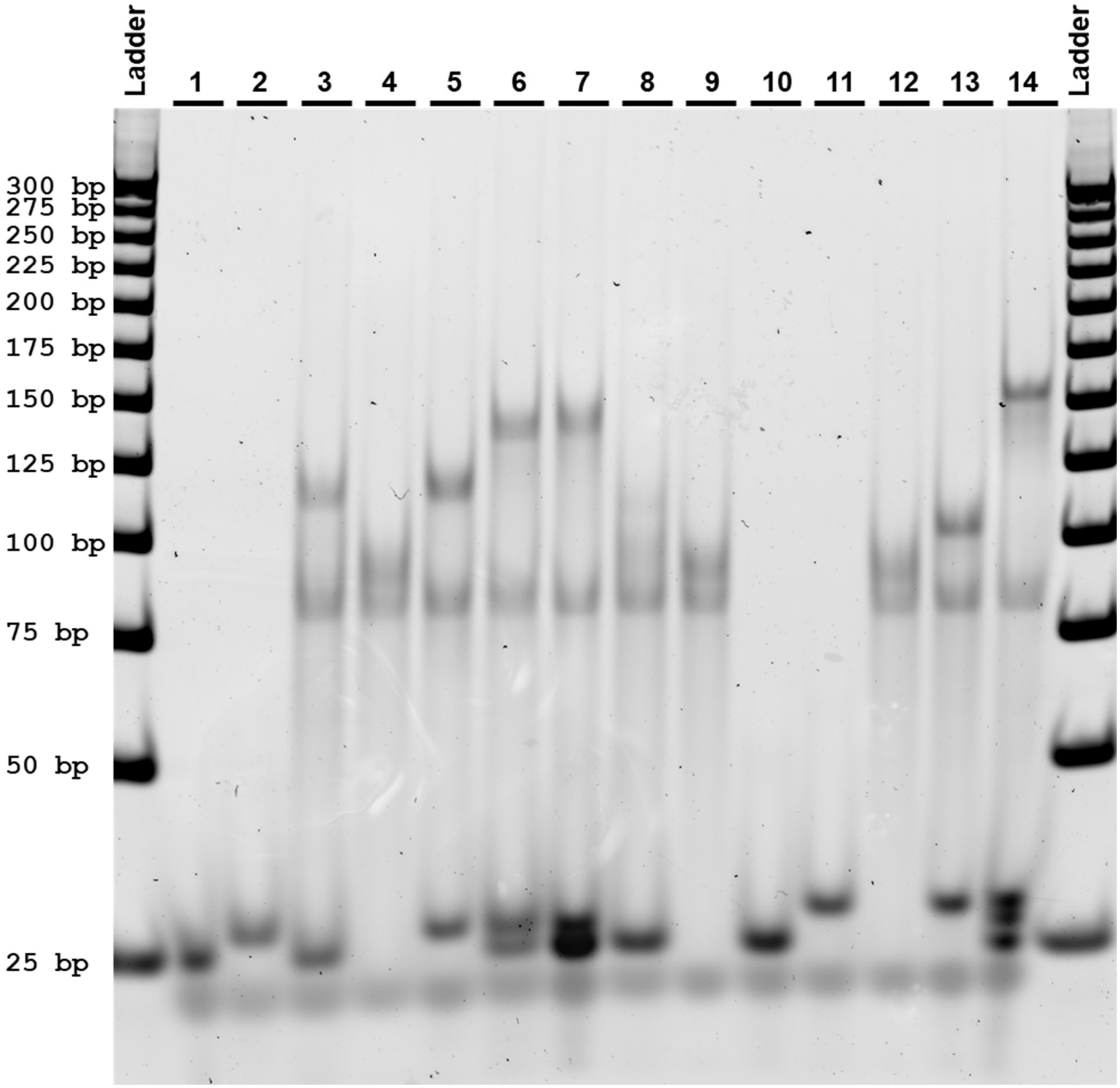


**Supplementary Fig. 11 | Native 5% PAGE assay of probe binding and target integrity after bisulfite treatment.** Lanes: 1, Aux1; 2, Aux2; 3, Aux1 + 102 nt MBC; 4, 102 nt MBC; 5, Aux2 + 102 nt MBC; 6, Aux1 + Aux2 + 102 nt MBC; 7, Aux1 + Aux2 + CP1 + 102 nt MBC; 8, CP1 + 102 nt MBC; 9, 102 nt MBC; 10, CP1; 11, CP2; 12, 102 nt MBC; 13, CP2 + 102 nt MBC; 14, Aux1 + Aux2 + CP2 + 102 nt MBC. Final concentration of all probes is 100 nM. MBC was diluted to 16 nM after quantification by Qubit. Before loading, all samples were incubated in 4× PBS with 2 µM dT10 in a thermocycler using the following protocol: denaturation at 73 °C or 80 °C for 3 min, annealing at 46.6 °C for 5 min and 40 °C and cooling down to RT (room temperature). In lanes 4, 9 and 12, both forward strand (top band) and backward strand (lower band) are observed on the gel, with migration between the 75 bp and 100 bp markers. In lane 8, a blurring band shows unstable binding of CP1 to MBC, suggesting possible degradation of the CP1 used in this experiment (see **Supplementary Fig. 2d,e**).
